# Human Rhinovirus 16 impacts cilia structure in 3D cultured primary bronchial epithelial tissue through alternative splicing of host cilia RNAs

**DOI:** 10.1101/2025.08.26.672366

**Authors:** Christoforos Rozario, Quan Gu, Andrew Stevenson, Rose A Maciewicz, Sheila V Graham

**Affiliations:** MRC-University of Glasgow Centre for Virus Research, School of Infection and Immunity, College of Medical, Veterinary and Life Sciences, University of Glasgow, Glasgow, G61 1QH, Scotland, UK; GLAZgo Discovery Centre, School of Infection and Immunity, College of Medical, Veterinary and Life Sciences, University of Glasgow, Glasgow, G12 8TT, Scotland, UK

**Keywords:** Human rhinovirus, respiratory epithelial cell, SR proteins, SRPK1, cilia

## Abstract

Human rhinoviruses (HRV) are a leading cause of the common cold but can often lead to respiratory complications such as wheeze in young children. In a transcriptomic study of respiratory nasal swab specimens from children hospitalised with acute wheeze, a significant alteration was found in the expression of the serine/arginine rich splicing factor (SRSF) kinase, SRPK1, between HRV positive children with acute exacerbations and HRV negative controls. As this kinase can regulate host RNA splicing, we hypothesised that HRV infection could dysregulate the expression of host mRNAs to affect antiviral functions or to alter the morphological features of the infected respiratory epithelium. Here, we show that pharmacological inhibition of SRPK1 in primary bronchial epithelial cells resulted in increased HRV16 replication while overexpression of SRPK1 reduced viral replication. In a primary bronchial epithelial 3D model infected with HRV16 decreased phosphorylation of SRSF1, 3 and 6 was observed. Furthermore, transcriptomic and alternative splicing (AS) bioinformatic analysis revealed the significantly altered AS of 1228 host genes during infection. Subsequent pathway analysis revealed the enrichment of most of these genes in networks related to cilia development and function. HRV16 infection led to significantly decreased cilia length and total cilia numbers in the primary bronchial epithelial 3D model together with changes to selected cilia proteins. Overall, this investigation has unravelled novel cellular networks implemented during HRV infection that may lead to acute exacerbations of respiratory infections.

**Author summary:** Human rhinoviruses cause the common cold. In immunocompetent individuals this is usually a self-limiting infection. However, in young children and the elderly, infection can lead to complications such as bronchiolitis, croup, and wheezing. Rhinovirus infection can exacerbate chronic conditions such as cystic fibrosis, chronic obstructive pulmonary disease and asthma. Understanding the molecular pathology of this exacerbation could lead to new avenues for therapy. In this study, we discovered that a multifunctional cellular enzyme called serine arginine protein kinase 1 (SRPK1) is a restriction factor for human rhinovirus 16 (HRV16) infection. One key cellular function of SRPK1 is to regulate RNA splicing through modifying the SR proteins that normally enhance splicing. In three dimensional tissues grown from human bronchial epithelial cells, we found that HRV16 infection led to decreased levels of modified SR proteins. This change resulted in significant alterations in RNA expression in the infected cell. Most of these alterations affected production of the correct versions of cilia proteins resulting in reduced cilia numbers and cilia blunting. This type of damage due to HRV infection would result in inefficient clearance of subsequent viral infections prolonging the viral infection leading to lower respiratory tract infection and to exacerbations of existing chronic disease.

## Introduction

Human rhinoviruses (HRVs) are positive-sense single stranded RNA enteroviruses within the Picornaviridae family, collectively comprising more than 160 genotypes grouped into 3 species (A, B and C) [1]. HRVs are placed amongst the most frequent human infectious agents, causing more than half of upper respiratory tract infections globally, though evidence now includes the lower respiratory tract in their niche [2]. They are causative agents of the common cold, a self-limiting infection with mild symptoms in immunocompetent individuals. However, HRV infection can lead to severe respiratory complications in immunocompromised groups such as pre-school children and the elderly. These complications are usually associated with the migration of the infection into the lower respiratory tract and include bronchiolitis, croup, wheezing as well as the exacerbation of chronic conditions such as cystic fibrosis, chronic obstructive pulmonary disease and asthma [2, 3].

Strong evidence associating HRV and asthma exacerbations has been accumulating, with HRV shown to account for more than 50% of total exacerbations and common cold complications in asthmatics costing about 60 billion USD annually [1]. Additionally, HRV-induced wheezing in early age is linked with asthma development in adulthood, while offspring of atopic (hyperallergic) mothers are more susceptible to HRV infections [4]. Despite all the evidence associating HRV with asthma, molecular mechanisms underlying this pathophysiology remain unclear.

The Mechanisms of Acute Viral Respiratory Infections in Children (MAVRIC) study conducted in Perth, Australia, aimed to investigate further the molecular mechanisms underlying asthma pathogenesis in the context of HRV infections. In this study, nasal swabs containing nasal epithelial cells along with a variety of immune cells including neutrophils and peripheral blood mononuclear cells were collected from pre-school children hospitalised with acute wheezing; these were subjected to microarray analysis comparing expression profiles between uninfected controls and HRV infected patients [5–8]. Surprisingly, for an RNA virus infection operating in the cytoplasm, genes involved in the pre-mRNA splicing process showed altered RNA expression.

It is well-established that HRV proteases 2A^pro^ and 3C^pro^ can degrade nuclear proteins and that proteases from the three different HRV species degrade different substrates [9]. Key substrates for HRV proteases are the nucleoporins that make up the nuclear pore complex (NPC) [9–11]. Degradation of nucleoporins results in mis-localisation of heterogenous ribonucleoprotein particles (hnRNPs) and SR splicing factors from the nucleus to the cytoplasm [12, 13]. It has been suggested that this virus-mediated relocation of splicing factors aids viral replication by supplying RNA genome-binding proteins [14] or by regulating virus mRNA translation [13, 15].

There are nine classical SR proteins (serine-arginine-rich splicing factors (SRSFs) 1-9). They all contain an N-terminal RNA recognition motif (RRM) and a C-terminal serine-arginine-rich (RS) domain [16]. Some SR proteins such as SRSF1 and SRSF6 possess an additional pseudo-RRM [17]. SRSFs are found predominantly in the nucleus but some (e.g. SRSF1) can dynamically shuttle to and from the cytoplasm [16]. SRSFs are essential regulators of constitutive and alternative splicing. They bind exonic or intronic sequence enhancers to define exon-intron boundaries and stabilise formation of the spliceosome at these boundaries to enhance splicing [18, 19]. A role for virus-associated splicing regulation in respiratory disease was strengthened when the SRSF6 gene was found to be upregulated in equine airway smooth muscle cells from asthmatic horses [20]. Several viruses of different Baltimore classification groups, for example, human papillomavirus [21], human immunodeficiency virus [22], influenza A virus [23], and alphavirus [24] have evolved to utilise or control SR protein family members and the host splicing machinery. SR proteins are also known to exert regulatory functions beyond splicing in the nucleus, including nuclear export, cytoplasmic stability and translation [25, 26].

SRPK1 is a moonlighting protein involved in numerous intracellular signalling pathways [27]. It has been shown to regulate innate immunity to viral infection [28]. However, it is a key regulator of cellular splicing. The RS domain of SR proteins is subject to phosphorylation by kinases including serine/arginine protein kinases (SRPK) and CDK-like kinases (Clk) [29, 30]. This post-translational modification regulates both the function and subcellular localisation of SRSFs [31, 32]. SRPK is normally present in the cytoplasm of cells where it phosphorylates newly synthesised SRSFs, to licence their entry into the nucleus [33]. There are three SRPKs in human cells (SRPK1, 2 and 3). Only SRPK1 and SRPK2 are expressed in epithelial cells. In the nucleus, SRPK1 interacts with Clk1 to promote splicing [30]. The dynamic phosphorylation of the SRSF RS domains by SRPK1 governs their levels, activity and cellular localisation [31, 34–37]. Not all SR proteins are equally affected by SRPK1 activity. For example, SR proteins e.g. SRSF1, with two RRMs may be phosphorylated to control splicing in a different manner from their single RRM–containing counterparts e.g. SRSF3 [38].

Here we show SRPK1 activity on SR proteins is reduced during HRV16 (a variant of HRV-A) infection of primary epithelial cells and that SRPK1 is a restriction factor for HRV16 infection. RNA-Seq analysis revealed significant host transcriptome changes between HRV16-infected and mock-infected 3D cultured primary bronchial epithelial cells similar to those found previously [39–43]. However, alternative splicing analysis of the RNA-Seq data revealed that splicing of RNAs involved in cilia structure and function was significantly altered. HRV16 infection impacted expression of key cilia components. This may indicate a sophisticated viral mechanism of host cell disruption during infection. The data suggest that HRV16 may regulate splicing of cilia-related RNAs leading to altered mucociliary clearance and ultimately prolonging productive viral replication.

## Materials and Methods

### Viruses stock generation

A variant of HRV-A, HRV16, was used for this study. Viral stocks obtained from ATCC (ATCC VR-283), were propagated in HeLa Ohio cells at 33°C, seeded at 70-80% confluence and grown in Dulbecco’s modified Eagle medium (DMEM) with 10% foetal bovine serum and 1% penicillin/streptomycin (Thermo Fisher Scientific, UK). Infected cell lysates were centrifuged at 120 x g for 7 min at 4°C. The supernatant was filtered through a 0.22 μM pore and stored at -80°C. Viral titres were determined via TCID_50_ assays using HeLa Ohio cells in 96-well plates. Cells were inoculated for 4 hours with serially diluted virus. Inoculates were washed off and cells were incubated for 5 days at 33°C. Cell death was recorded, with each dilution tested in quadruplicate and viral titre calculated using the Spearman & Karber method [44].

### Cell growth

Normal Human Bronchial Epithelial cells (HBECs) were purchased from Lonza (Basel, Switzerland # CC-2540S). The cell donor was a female Caucasian aged 16 years old, with no known disease or smoking history and a BMI of 22. Cells were grown in PneumaCult-Ex plus medium (Stem Cell Technologies, Cambridge, UK) in T75 flasks for 2D culture. For air-liquid interface 3D culture, cells were seeded on 6.5 mm transwells with 0.4 µm pore polyester membranes (Corning, Berlin, Germany). When cells reached full confluence, airlift was performed by aspirating the apical medium and replacing the basal medium with PneumaCult ALI maintenance medium (Stem Cell Technologies, Cambridge, UK). Medium was replaced 3 times per week, with initial mucus production at approximately 2 weeks post-airlift. Mucus was washed off once per week and tissues reached full differentiation by 4 weeks post-airlift.

### HBEC infections

A mucus wash and basolateral medium change were performed. Cultures were incubated for one hour at 33°C while virus stocks were thawed on ice. Based on calculations of an *in vivo* infectious dose [40] the inoculum (3×10^6^ pfu) was added in 100 µl medium apically. The same volume of medium was added to the top of mock-infected control cultures. Tissues were incubated for 3.5 hours at 33°C then the inoculum was aspirated off and tissues were washed apically three times with PBS without Ca^2+^ and Mg^2+^ to remove non-internalised virus. Tissues were then incubated 33°C for the times stated in the experiments.

### SRPK1 overexpression, depletion and SRPIN340 inhibition

The SRPK1 (transcript variant 1) human cDNA clone (untagged) (OriGene,Herford, Germany, #RC205315) was transfected using Lipofectamine2000 (Thermo Fisher Scientific, UK) at a concentration of 200ng/ml for 48 hours. SRPK1 was depleted by transfecting Dharmacon SMART-Pool siRNAs in RNAiMax transfection reagent (Thermo Fisher Scientific, UK) into bronchial epithelial cells. siGLO (Dharmacon, #<u>D-001630-01</u>) was used as a non-target siRNA control and to monitor transfection efficiency. SRPIN340 (Sigma Aldrich, UK, #5.04293) was dissolved in DMSO to 20mM stocks and was administered at 20μM for 48 hours. Both treatments were administered to primary bronchial epithelial cells seeded at 60% confluence.

### RNA extraction and RT-qPCR

2D cultures: Cells grown in 6-well plates were lysed in 500 μl of Trizol reagent (Thermo Fisher Scientific, UK) and stored at -20°C. Upon thawing cells were scraped into Trizol and RNA was isolated according to the manufacturer’s instructions. 3D cultures: Tissues were flash frozen in liquid nitrogen and stored at -80°C. Upon thawing cells were vortexed to detach from the transwell membrane then tissues were ground to a fine powder under liquid nitrogen using a mortar and pestle and RNA was isolated using the RNeasy extraction kit (Qiagen, Germany) according to the manufacturer’s instructions. RNA was quantified using a NanoDrop 2000 Spectrophotometer (Thermo Fisher Scientific, UK, #ND-2000).

cDNAs were synthesised using the Maxima First Strand cDNA Synthesis kit with DNase digestion according to the manufacturer’s instructions (Thermo Fisher Scientific, UK). 20 μl reactions were prepared using the Takyon™ ROX Probe 2X MasterMix dTTP blue (Eurogentec, Camberly, UK), primers and probes at 300 and 100 nM respectively and nuclease-free water. Primer/probe sets used are listed in Table 1.

**Table 1.**
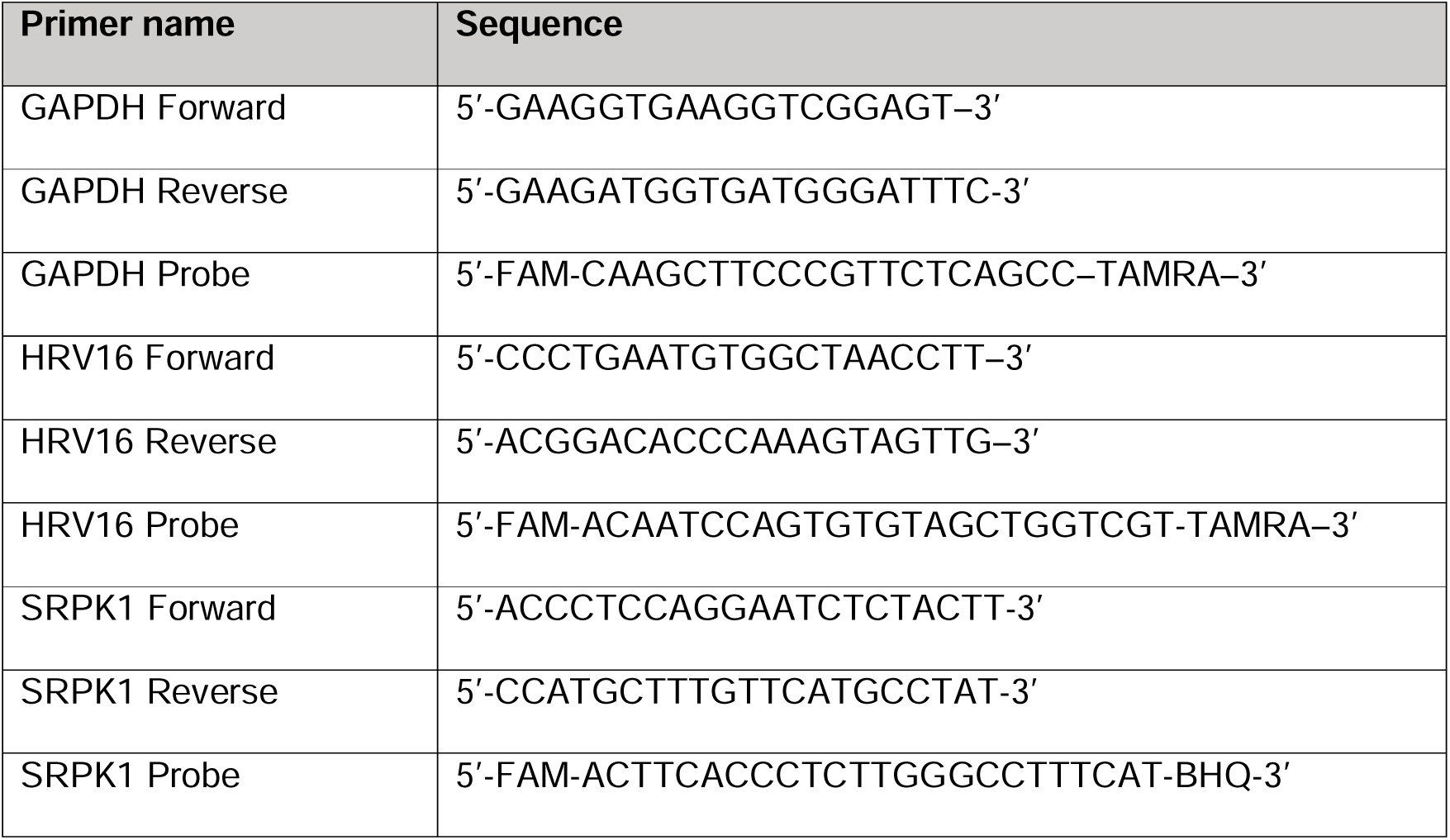
List of primers and probes used in RT-qPCR experiments.

Reactions were run on an ABI 7500 thermocycler (Thermo Fisher Scientific, UK) at this profile: 95°C (5 min), 60°C (15 sec), 72°C (3 min), 40 cycles. Data analysis was performed using the 7500 v2.3 (Thermo Fisher Scientific, UK) software. Ct values were determined relative to GAPDH as the reference target gene.

### Protein extraction and western blotting

Cells grown in 6-well plates were scraped in 400 μl 2X Bolt LDS buffer (Thermo Fisher Scientific) containing PhosphoSTOP (Merck, UK catalogue # 04693116001) and complete miniprotease inhibitor cocktail (Merck, UK, catalogue # 200-664-3) in PBS. 20 µl of sample was loaded per lane on Bolt 4-12% Bis-Tris polyacrylamide gels (Thermo Fisher Scientific, UK) and electrophoresed at 150 V for 60 minutes. Proteins were transferred to nitrocellulose membranes using the iBlot2 Dry Blotting System (Thermo Fisher Scientific, UK). Membranes were blocked in 5% (w/v) milk powder in PBS containing 0.01% (v/v) Tween. PBST at room temperature for 1 hour, then washed in PBST (3 x 7 minutes) and incubated with primary antibody in 5% milk powder in PBST for 1 hour at room temperature or overnight at 4°C with rotation. Following incubation, membranes were washed in PBST (3 x 7 minutes) and incubated with secondary antibody in 5% milk powder in PBST for 1 hour at room temperature in the dark. A final round of washes was performed in PBST (2 x 7 minutes) and PBS (2 x 7 minutes), and bands were visualised on the LI-COR Odyssey CLx Infrared imaging system. Primary antibodies were SRPK1 1:300 (1:500, clone G211-637 BD Transduction Laboratories, catalogue #611072), SRSF1 1:2500 1:1000, Mab96, Thermo Fisher Scientific, catalogue # 32-4500), SRSF3 (1:300, Life Technologies, UK, catalogue #334200), SRSF6 (1:300 Abcam, UK, catalogue #ab140623, GAPDH (1:1000, Meridian Life Sciences, UK, catalogue #H86504M, clone 6C5), HRV16 VP0/VP2 (1:250, QED Biosciences, Aachen, Germany, catalogue #<u>18758</u>). mAb104-detecting phosphorylated SRSFs was prepared from hybridoma (ATCC CRL-2067) supernatants and was used neat with 5% milk powder and 0.01% Tween. Secondary antibodies were goat anti-rabbit Dylight 800 conjugate (1:2000, Thermo-Fisher, UK, catalogue #SA5-35571), goat anti-mouse Dylight 800 conjugate (1:2000, Thermo-Fisher, UK, catalogue #SA5-35521) and IRDye anti-mouse 800CW (1:2000, IRDye Licor Biosciences Ltd, UK, catalogue #926-32210). Membranes were imaged on an Odyssey Infrared Imager (LiCOR). The intensity of protein bands was quantified using Odyssey Image Studio software. Protein levels were determined and normalised to the level of the endogenous control (GAPDH).

### Formalin fixed paraffin embedded (FFPE) sample preparation

Primary bronchial epithelial 3D cultures were fixed by fully submerging in 10% (v/v) buffered formaldehyde (BNF) at room temperature overnight. The cultures were submitted to the Veterinary Diagnostic Services (University of Glasgow) for paraffin embedding, and haematoxylin and eosin (H&E) staining.

### Immunofluorescence microscopy

Antigen retrieval for 4µm sections from formalin-fixed, paraffin-embedded samples was carried out in 10 mM sodium citrate buffer, pH 6.0 using a Menarini Access Retrieval Unit, at 110^0^C on full pressure for 10min. (Veterinary Diagnostic Services, University of Glasgow). Microscope slides were washed sequentially six times in PBST. Slides were blocked in 10% (v/v) filtered donkey serum in PBS for 1 hour at room temperature. Sections were incubated with primary antibody diluted in 5% donkey serum in PBS for 2 hours at room temperature. Slides were washed and sections were incubated with secondary antibody diluted in 5% donkey serum in PBS for 2 hours at room temperature under dark conditions. Slides were mounted in ProLong Gold Antifade Mountant with DAPI (Thermo Fisher, UK, catalogue # P36931) and visualised on a ZEISS LSM 710 confocal microscope. Images were acquired using the ZEN Blue software. Primary antibodies were HRV16 VP0/VP2 (1:100, QED Biosciences, Aachen, Germany, catalogue #<u>18758</u>), β-tubulin (1:250,Merck, UK, catalogue #AB9354) and TMEM67 (1:250, Proteintech, UK, catalogue #Ag5009). Secondary antibody: donkey anti-mouse Alexa-fluor 555-labelled antibody (1:1000 Thermo Fisher Scientific, UK, catalogue #A-31570).

### RNA sequencing, differential expression and pathway analysis

3D bronchial epithelial cultures were infected (3×10^6^ pfu) at four weeks post-airlift and harvested 48 hours post-infection. Total RNA was prepared from 3 biological replicates per condition and sequenced in-house. Eluted RNA was quantified using a NanoDrop 2000 Spectrophotometer (Thermo Fisher Scientific, ND-2000) and quality controlled on a TapeStation (Agilent Technologies, G2991AA). All samples had a RIN score of ≥ 9. One microgram of total RNA was used to prepare libraries for sequencing using an Illumina TruSeq Stranded mRNA HT kit (Illumina, #20020594) and SuperScript2 Reverse Transcriptase (Thermo Fisher Scientific, #18064014) according to the manufacturer’s instructions. Libraries were pooled in equimolar concentrations and sequenced using an Illumina NextSeq 500 sequencer (Illumina, #FC-404). RNA-Seq reads were analysed for quality using FastQC (version 0.11.9) and reads were trimmed of adaptor sequences and low-quality bases using Trimgalore (https://www.bioinformatics.babraham.ac.uk/projects/trim_galore/). The trimmed reads were aligned to the human genome GRCh38 (*Ensembl*) using Hisat2 (version 2.2.0) [45]. FeatureCounts (Version 2.0.1) [46] was used to quantify reads mapping to gene annotation files. Reads counts were normalized to counts per million (CPM). The edgeR package was used to calculate the gene expression level and to analyse differentially expressed genes between sample groups. RNA-Seq data sets are freely available from the European Nucleotide Archive accession number PRJEB88791, and heat maps were generated in GraphPad Prism (version 9). Pathway analysis was performed using the Ingenuity Pathway Analysis (IPA) tool using the rat, mouse, human and undefined species data, in all cells and tissues. Only experimentally observed data were selected.

### Over representation analysis of alternative splicing events

Bam files from the RNA paired-end sequencing were sorted by co-ordinate, indexed and subject to SplAdder analysis [47] to measure and quantify alternative splicing events. Percentage spliced in (PSI) values were quantified for each splicing event, and a two-tailed student’s t-test was performed for values from mock-infected and HRV16-infected primary bronchial epithelial cells to determine the most significantly differentially spliced genes. Pathway analysis was performed using Webgestalt (http://www.webgestalt.org/) [48]. Over representation analysis was carried out test for biological processes using Benjamini-Hochberg multiple testing adjustment. Sashimi plots were generated using MISO (https://pypi.org/project/misopy/0.5.4/) [49].

### Cilia count

Five sections from three biological replicates per condition were randomly selected and cilia were manually counted from images obtained at x20 magnification from two technical replicates per condition using a tally meter from FFPE H&E-stained samples. Image J ( https://imagej.net/software/fiji) was used to quantify the cilia length by measuring the proportion of distance in pixels. Ten cilia were measured in each section for each condition.

### Statistical analysis

Statistical analyses were carried out using a students’ t-test. P-values of < 0.05 were considered statistically significant.

## Results

### SRPK1 restricts HRV16 infection in primary epithelial cell culture

The kinase SRPK1 was found to be up-regulated at the mRNA level in the MAVRIC study which compared nasal swab samples from HRV-infected young children with wheezing exacerbations to HRV-negative controls to [7, 50]. However, when the study population was divided into two different phenotypes based on a Th1/type 1 interferon response versus a Th2/IFNγ response, SRPK1 mRNA was significantly downregulated in the second population. The two phenotypes had quite different clinical characteristics. In the case of a Th2/IFNγ response, illness progressed more slowly but there was a greater chance of hospitalisation and repeat infections/exacerbation of disease [7] suggesting that SRPK1 downregulation was associated with more severe disease.

However, SRPK1 activity, as opposed to SRPK1 levels, is regulated by key cell signalling pathways such as CK2 and Akt, which can be impacted upon virus infection [51]. To find out more about the relationship between HRV infection and SRPK1, we overexpressed the kinase in human bronchial epithelial cells (HBECs). We also inhibited the kinase by treating cells with 20µM SRPIN340, a specific inhibitor of the kinase activity of SRPK1 [52]. Each treatment was carried out 48 hours prior to HRV16 infection. We examined changes due to the treatments in the logarithmic phase of viral production (MOI=3, 16 hours post infection, Supplementary Fig. 1A). Supplementary Fig. 1B shows a significant increase in SRPK1 mRNA levels in overexpressing cells relative to mock-transfected cells. The effectiveness of the SRPIN340 treatment was shown through decreased phosphorylation of SRSF1 (Supplementary Fig. 1C) and SRSF6 (Supplementary Fig. 1E) compared to mock-treated cells. Levels of phosphorylated SRSF3 were not significantly decreased by SRPIN340 treatment (Supplementary Fig. 1D). No significant change in levels of phosphorylated SRSFs was detected when SRPK1 was overexpressed (SRPK1 OE) (Supplementary Fig. 1C-E).

We assessed how inhibition and overexpression of SRPK1 impacted HRV16 infection. The levels of viral RNA, tissue-released virus particles and expression of viral capsid proteins VP0 and VP2 were compared between mock-treated, SRPIN340-treated and SRPK1-overexpressed HBECs infected with HRV16 for 16 hours at an MOI=3 (Fig. 2). SRPK1 inhibition led to increased HRV16 replication, as reflected by the increased viral RNA production (Fig. 2A, SRPIN340) although this was not statistically significant (p=0.07). However, a statistically significant increase was observed upon kinase inhibition in tissue-released virus particles (p<0.05) (Fig. 2B, SRPIN340) and virus capsid VP2 protein production (Fig. 2C, D, SRPIN340). In contrast, SRPK1 overexpression did not have a significant effect on viral genome replication when compared to untreated controls (Fig. 2A, SRPK1 OE). SRPK1-overexpressing HBECs showed significantly decreased virus shedding (Fig. 2B, SRPK1 OE) and viral capsid protein VP0 production (Fig 2C, D, SRPK1 OE) compared to untreated samples, thus displaying an opposite effect from respective SRPK1-inhibited samples. Taken together, these data suggest that SRPK1 is a restriction factor for HRV16 infection in primary epithelial cells, with the activity of SRPK1 repressing viral replication, assembly and release, at 16 hours post-infection.

**Figure 1.**
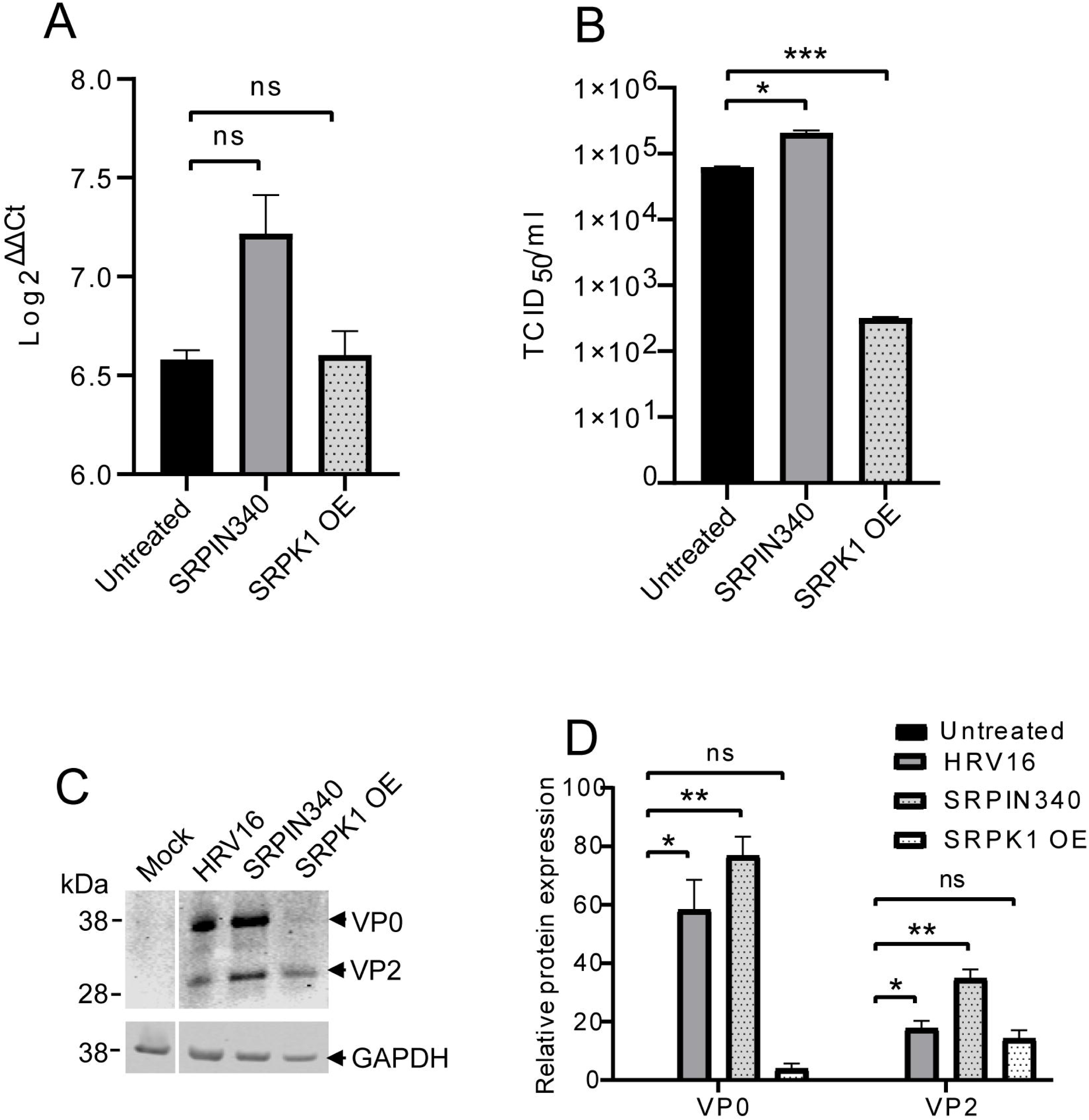
Changes in SRPK1 levels and activity alter HRV16 infection in primary human bronchial epithelial cells. A. RT-qPCR analysis of tissue-associated HRV16 RNA levels in untreated, SRPIN340-treated and (SRPIN340) SRPK1 overexpressing (SRPK1 OE) HBECs infected with HRV16 (MOI=3) for 16 hours. The Log2^ΔΔCt^ values were calculated using the values of the housekeeping gene GAPDH as the control target. B. TCID_50_ quantification of tissue-released infective virus using supernatants from untreated, SRPIN340-treated (SRPIN340) and SRPK1 overexpressing (SRPK1 OE) HBECs infected with HRV16 (MOI=3) for 48 hours. C. Western blot analysis of the HRV16 capsid proteins VP0/VP2 expression in mock-treated and mock-treated SRPIN340-treated or SRPK1 overexpressed (OE) cells infected with HRV16 (MOI=3) for 48 hours. A GAPDH loading control is shown under the virus capsid protein blot. D. Quantification of western blot data from three separate experiments. All data show the mean and standard error from three separate experiments. ns=non-significant statistical difference, *=p<0.05, **=p<0.005, ***=p<0.0005 (student’s t-test, 2-tailed).

**Figure 2.**
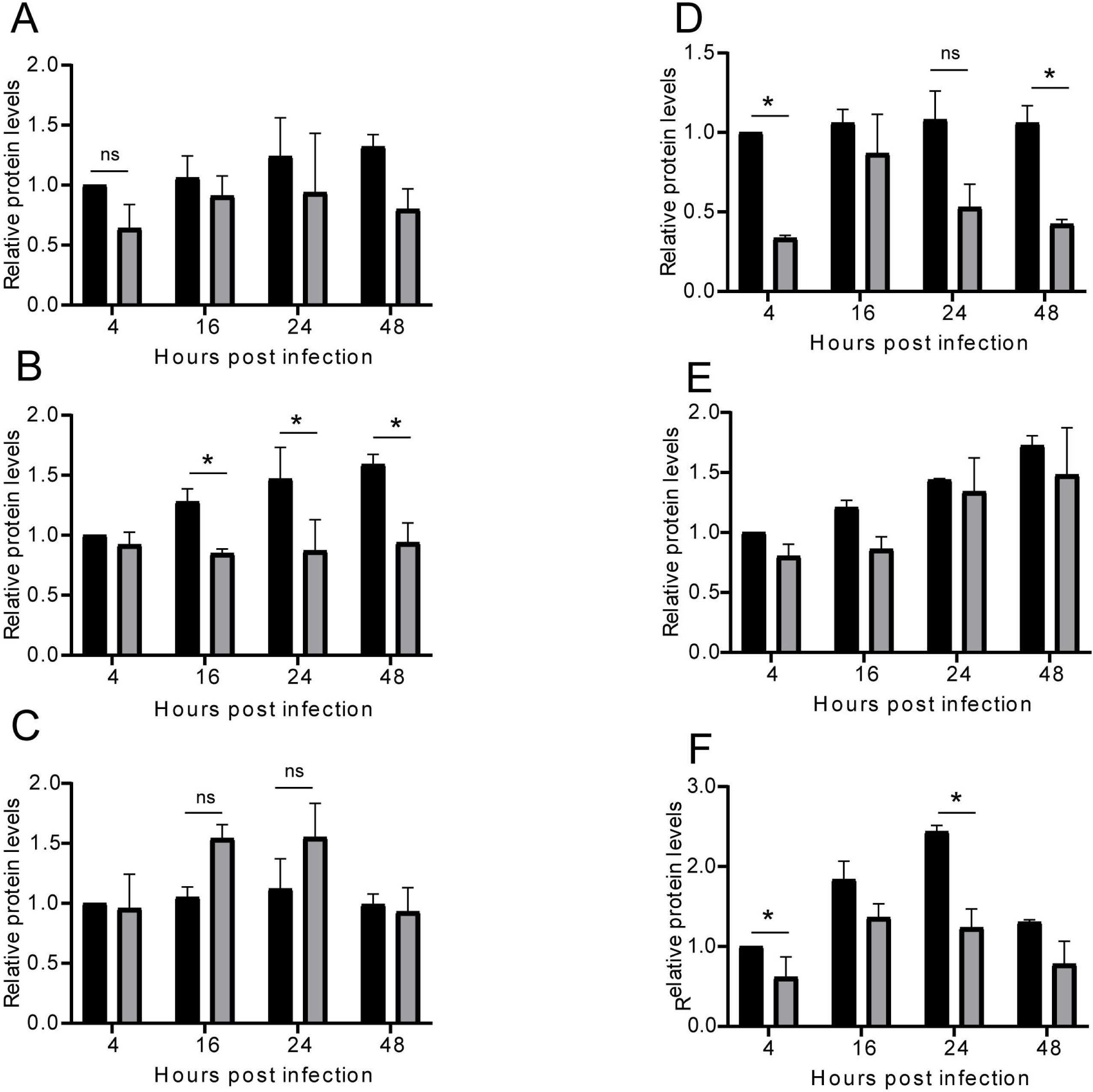
HRV16 infection of primary human bronchial epithelial cells downregulates expression of SRSF3 and decreases phosphorylation of SRSF1 and SRSF6. Quantification of western blot expression data relative to GAPDH expression of A. total SRSF1, B. total SRSF3, C. total SRSF6, D. phosphorylated SRSF1, E. phosphorylated SRSF3, and F. phosphorylated SRSF6 in mock-infected (Control) and HRV16-infected (HRV16: MOI=3) HBECs at 4, 16, 24 and 48 hours post-infection. The data show the mean and standard error of the mean from three separate experiments. Significant statistical difference, *=p<0.05, **=p<0.005, ***=p<0.0005 (student’s t-test, 2-tailed).

### HRV16 infection alters phosphorylation of SR proteins in primary human bronchial epithelial cells

Next, to assess activity of SRPK1 during HRV16 infection we quantified changes in expression of selected substrates of SRPK1 in HBECs. SRSF1, SRSF3 and SRSF6 were selected for further study. SRSF1 is the prototypical SRSF protein [37, 53]. SRSF3 was previously shown to be involved with the internal ribosome entry site (IRES)-dependent translation of picornavirus mRNAs [54] and SRSF6 is known to be upregulated in equine airway smooth muscle cells from asthmatic horses [20]. There was no significant change in total levels of SRSF1 (Fig. 2A, Supplementary Fig. 2A) or SRSF6 (Fig 2C, Supplementary Fig. 2A) during a 48 hour time course of infection (MOI=3). However, SRSF3 showed significantly decreased expression at 16, 24 and 48 hours of infection (Fig. 2B, Supplementary Fig. 2A). Next, we measured levels of phosphorylated SR proteins using the SR protein phosphor-specific antibody Mab104 [38]. Levels of phosphorylated SRSF1 were significantly decreased at 4 and 48 hours post infection in HRV16-infected cells compared to mock-infected cells (Fig. 2D, Supplementary Fig. 2B). There was no significant change in levels of phosphorylated SRSF3 at any time point in HRV16-infected cells compared to mock infected cells (Fig. 2E, Supplementary Fig. 2B). Finally, phosphorylated SRSF6 levels were significantly decreased in HRV16-infected cells at 4 and 24 hours post infection (Fig. 2F, Supplementary Fig. 2B). Taken together, these data indicate that HRV16 infection represses the expression of SRSF3 and inhibits phosphorylation of SRSF1 and SRSF6. The downregulated phosphorylation seen for SRSF1 and SRSF6 suggests that the activity of SRPK1 is repressed at early and late times of infection during HRV16 infection.

### HRV16 infection in 3D cultured primary bronchial epithelial tissues inhibits SRPK1 protein levels and phosphorylation of SR proteins

Next, we evaluated levels and activity of SRPK1 in HRV16 infection of air-liquid interface 3D tissue cultures of HBECs. 3D cultures enable the differentiation of these cells, providing increased physiological relevance in parameters such as cell polarisation, cilia formation, cell to cell interactions and nutrient access [55]. HBECs were grown at the air-liquid interface in transwell cultures over a four-week period (Supplementary Fig. 3 compare A to B). After this time, the tissues were fully differentiated as indicated by cilia development (Supplementary Fig. 3B, arrowhead) and a multi-layered pseudostratified epithelium was developed. The tissues also produced mucus as indicated by periodic acid-Schiff (PAS) staining of the tissues which detects polysaccharides (Supplementary Fig. 3, compare C to D). This suggests the presence of goblet cells (Supplementary Fig. 3D, arrowheads) further confirming tissue differentiation. HRV16 (3×10^6^ pfu) was applied to the surface of the epithelium and infection was allowed to proceed for 48 hours. Infection induced evident mechanical damage to the structure of the epithelium (Supplementary Fig. 3 compare E and F, arrows in F shows regions of damage). However, the extent of damage may not be due only to HRV16 infection but also due to tissue handling for preparation of formalin-fixed paraffin-embedded blocks. Immunofluorescence staining to detect the virus capsid protein VP2 indicated that infection was established in all the layers of the epithelium (Supplementary Fig. 3 compare G and H). Western blot analysis indicated a significant decrease in SRPK1 and phosphorylated SRSF1, SRSF3, and SRSF6 protein levels in 3D HBEC tissues infected with HRV16 (3×10^6^ pfu) for 48 hours compare to mock-infected tissues (Fig. 3).

**Figure 3.**
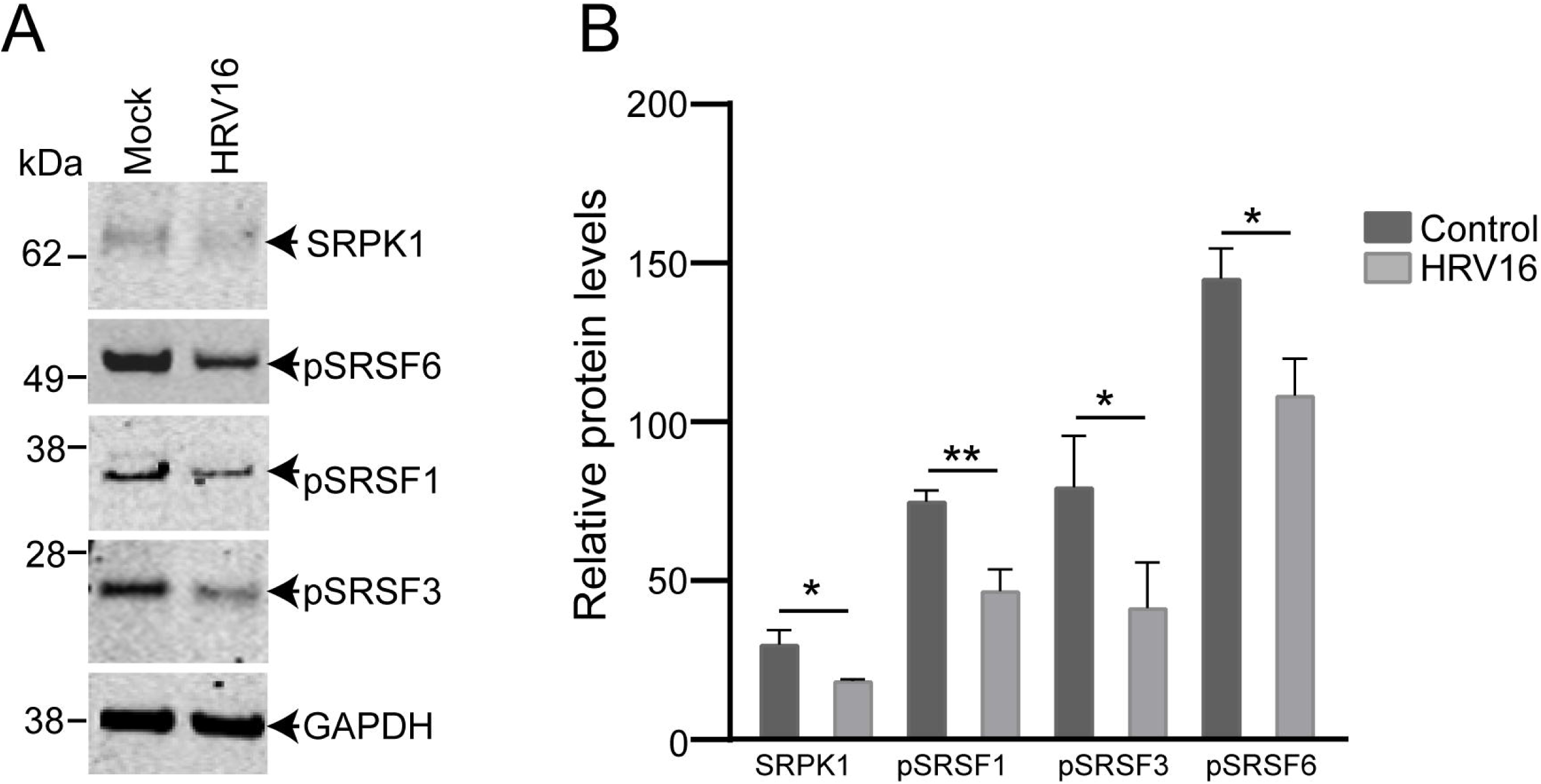
HRV16 infection in 3D cultured primary bronchial epithelial tissues significantly decreases the phosphorylation of SRSF proteins. A. Western blot analysis of levels of SRPK1 and phosphorylated SRSF1, SRSF3, and SRSF6 (detected with Mab104) in mock-infected (Mock) and HRV16-infected (HRV16) HBEC air-liquid interface cultures at 48 hours post-infection (hpi) with 3×10^6^ pfu HRV16. GAPDH is shown as a loading control and was used to determine relative proteins levels. B. Quantification of data from three separate experiments. The data show the mean and standard error of the mean from three separate experiments. ns=no significant statistical difference. Significant statistical difference, *p<0.05, **p<0.005 (student’s t-test, 2-tailed).

### Transcriptomic analysis confirms increased expression of innate immune genes during HRV16 infection of 3D cultured primary bronchial epithelial tissues

RNA sequencing was performed on HRV16 and mock infected primary bronchial epithelial 3D cultured tissues. Tissues were mock-infected or infected (3×10^6^ pfu) for 48 hours at 4 weeks post-airlifting. Total RNA extracts were prepared from triplicate HRV16-infected and mock-infected HBEC 3D air-liquid interface cultures. The counts per million (CPM) of individual transcripts were similar between infected (median=33.1 CPM per transcript) and mock infected tissues (median=30.2 CPM per transcript). 82.2% of bases achieved a quality score of Q30 and Illumina software was used to assign sequencing reads to their corresponding samples. Differentially expressed genes (DEGs) between infected and mock-infected samples were identified by mapping sequencing reads to the reference human genome and quantifying the mapped reads of individual transcripts.

Analysis indicated that there were 4034 DEGs due to HRV16 infection (false discovery rate (FDR)<0.05, Benjamini-Hochberg correction). 2143 genes were up-regulated, and 1891 genes were down-regulated in infected when compared to mock-infected tissues (Supplementary Table 1). The most statistically significant differences in gene expression during infection were seen in up-regulated genes as illustrated in the volcano plot in Fig. 4A (red dots). Key innate immune gene changes are indicated similar to those found by Ong et al [43]. The violin plot in Fig. 4B shows that the distribution of gene expression was similar between infected and mock-infected samples. However, more genes showed a higher level of expression (increased reads per million) in control mock-infected (Ctrs) compared to infected (HRV16) samples (Fig. 4B).

**Figure 4.**
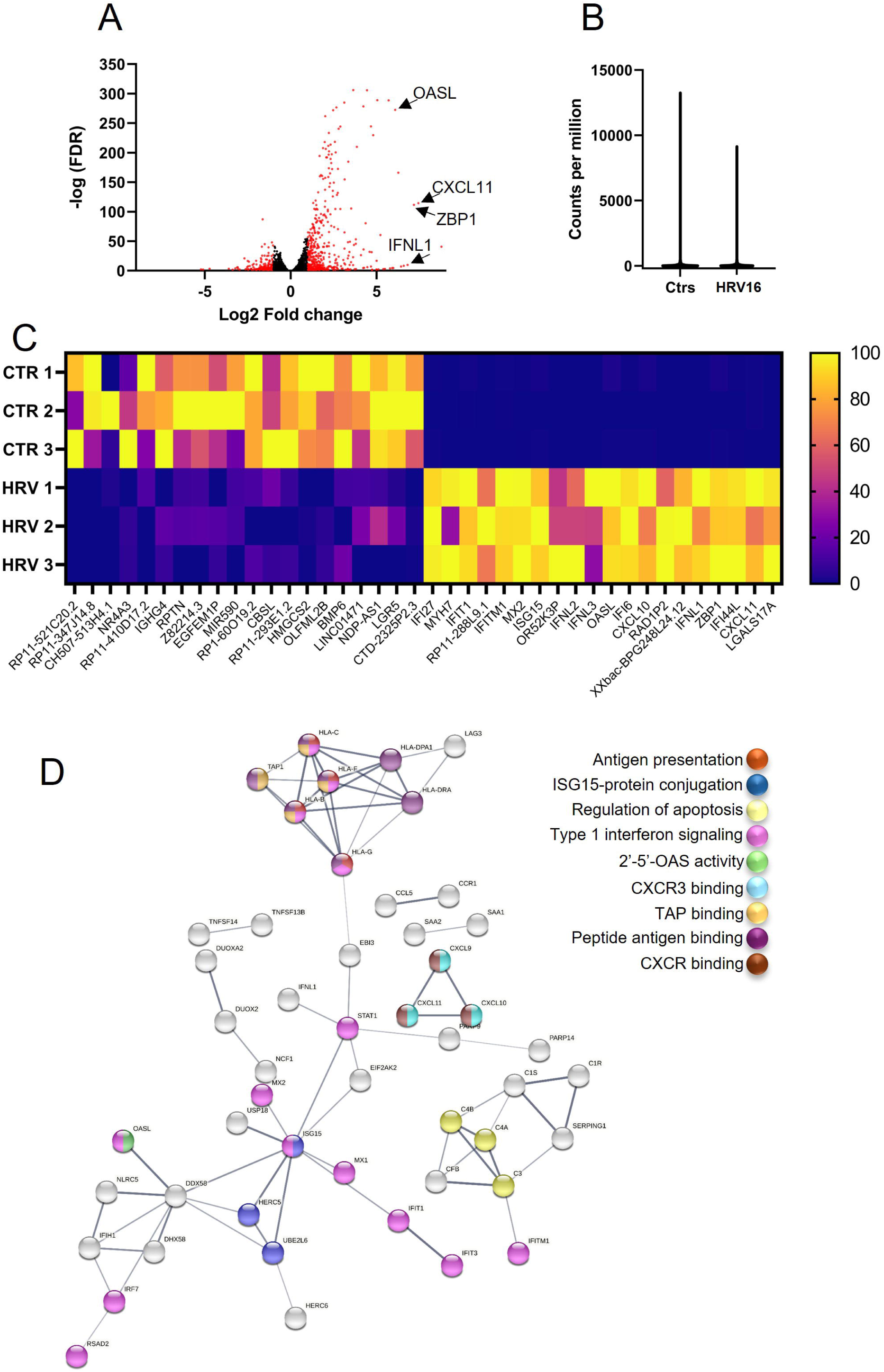
HRV16 induced changes in gene expression during infection in 3D cultured primary bronchial epithelial tissues. 3D tissues were cultured at the air-liquid interface for 4 weeks, then mock infected or infected apically with 3×10^6^ pfu HRV16 for 48 hours. RNA sequencing was performed on three replicates for each condition. A. Volcano plot of differentially expressed genes (DEGs) (scattered dots, n=4034) between infected and mock-infected samples (FDR<0.05). The x-axis is the log2 fold change (infected/controls) while the y-axis is the -log(FDR) calculated value. Black dots indicate the DEGs with log2 fold change within the range of -1 to 1. Red dots indicate DEGs with changes less than -1 or greater than 1. B. Violin plot comparing the distribution of gene expression between mock-infected (Ctrs) and infected (HRV16) tissues. C. Heatmap of the top 20 upregulated and downregulated genes in infected (HRV) and mock-infected (CTR) samples. Each row represents 1 of 3 replicates per condition. The color scale ranges from deep purple (no expression) to yellow (high level expression). D. Gene networks induced during HRV16 infection of HBECs in air-liquid interface culture. PPI network functional enrichment analysis (generated using STRING) using DEGs with log2-fold changes of greater than 2 or less than -2. Nodes represent the interacting proteins with lines representing direct links. A colour key shows the different networks. Up to four colours per node are shown.

Fig. 4C shows a heat map of the top 20 genes that were up- or down-regulated due to HRV16 infection. The most up-regulated genes were involved in anti-viral defence, primarily the type I interferon signalling pathway (Fig. 2A arrows, Fig. 4C, Table 2), demonstrated by the strong up-regulation of these genes in HRV16-infected samples, across all biological replicates as previously reported [39, 43, 56, 57]. On the other hand, the functions of the most down-regulated genes during infection were very variable, but included genes involved in epithelial homeostasis, metabolic processes, and cell signalling (Fig. 4C). Although SRPK1 was significantly downregulated at the protein level in HRV16 infected primary bronchial epithelial 3D cultured tissues (Fig. 3), SRPK1 mRNA expression was found to be significantly upregulated by 1.3-fold. Protein-protein interaction networks were constructed using Gene Ontology (GO) enrichment analysis of up-(240) or down-regulated (9) DEGs with log2 fold change >2 or <-2 (Fig. 4D). The antigen presentation pathway was the most enriched biological process. ISG15 protein conjugation, apoptosis and type 1 interferon signalling were also identified as biological pathways induced by HRV16 infection. Molecular function enrichment indicated HRV16-induction of antiviral OAS activity, chemokine receptors, and antigen processing (Fig. 4D).

**Table 2.** List of top 30 immune-related genes up-regulated by HRV16 infection.

| Log2-fold change | Gene name | Gene function |
| --- | --- | --- |
| 7.434 | CXCL11 | Chemokine |
| 7.434 | ZBP1 | Innate sensor |
| 7.223 | IFNL1 | Antiviral response |
| 6.268 | CXCL10 | Chemokine |
| 6.101 | IFI6 | Induced by interferon |
| 6.085 | OASL | Innate immunity to viral infection |
| 5.971 | IFNL3 | Antiviral response |
| 5.912 | IFNL2 | Antiviral response |
| 5.705 | ISG15 | Innate immunity to viral infection |
| 5.631 | MX2 | Innate immunity to viral infection |
| 5.559 | IFITM1 | Induced by interferon |
| 5.423 | IFIT1 | Induced by interferon |
| 5.239 | IFI27 | Induced by interferon |
| 5.229 | AIM2 | Innate immunity to viral infection |
| 5.049 | IFIT3 | Induced by interferon |
| 5.043 | EPSTI1 | Induced by interferon |
| 5.004 | CXCL9 | Chemokine |
| 4.810 | HERC5 | Innate immunity to viral infection |
| 4.672 | DUOXA2 | Antiviral protein in respiratory epithelial cells |
| 4.447 | CMPK2 | Innate immunity to viral infection |
| 4.370 | CSF3 | Controls production of granulocytes |
| 4.303 | RSAD2 | Antiviral protein |
| 4.257 | OAS3 | Innate immunity to viral infection |
| 4.238 | XAF1 | Induced by interferon |
| 4.141 | MX1 | Innate immunity to viral infection |
| 4.121 | BST2 | Innate immunity to viral infection |
| 4.007 | HERC6 | Innate immunity to viral infection |
| 3.938 | OAS2 | Innate immunity to viral infection |
| 3.796 | HLA-F | Major histocompatibility protein |
| 3.768 | IFI44 | Induced by interferon |
| 3.404 | IFITM3 | Induced by interferon |

Over-representation analysis was performed to identify cellular pathways affected by HRV16 infection. A subset of DEGs was selected for this pathway analysis, with log2-fold change values within the range of 1 to -1. This cut-off point included 655 DEGs of which 466 were up-regulated and 189 down-regulated during infection (Supplementary Table 2). Pathway analysis mapped the vast majority of the DEGs in innate immune pathways, with the interferon signalling pathway scoring the highest enrichment value (Supplementary Fig. 4). However, genes involved in epithelial cell junctions (claudins and cadherins) and the epithelial barrier (repetin, involucrin) were also downregulated as expected [58] (Supplementary Table 2). Table 2 lists the top thirty upregulated protein-coding genes of known antiviral functions.

### HRV16 infection of 3D cultured primary bronchial epithelial tissues leads to changes in the splicing of cilia-related genes

Since we had shown that HRV16 infection alters splicing factors, we wanted to assess whether the splicing of host pre-mRNA transcripts was altered during infection. We used the alternative splicing toolbox SplAdder [47] to perform further analysis on our RNA-seq data. Using SplAdder, we identified the total number of single splicing events in our dataset, along with the alternative splicing (AS) mechanism of each particular event. We identified 10952 genes with alternative splicing events, with each event categorized by its splicing mechanism (e.g., exon skipping, intron retention etc.). The differential splicing between infected and mock-infected samples was determined by comparing the percentage spliced-in (PSI) output values given for each transcript. PSI indicates the efficiency of splicing of a particular exon into the transcript population of a gene, with values ranging from 0 to 1. Therefore, different PSI values for a given transcript between infected and mock-infected samples, indicate a different AS pattern for that respective gene during infection. 10514 out of 10952 genes (96%) had a different PSI value in infected compared to mock-infected samples. Following a 2-tailed student’s t-test analysis, this number was limited to 1228 genes that were significantly differentially spliced due to HRV16 infection (11% of all the alternatively spliced genes in the dataset).

1469 single splicing events specifically associated with HRV16 infection were identified as being involved in producing mRNAs as some were alternatively spliced via more than a single mechanism, of which about 40% were exon skipping events (Fig. 5A). The rest of the splicing events identified included alternative 3’ splice sites (23%), alternative 5’ splice sites (18%), intron retention (10%), multiple exon skipping (8%) and mutually exclusive exons (2%) (Fig. 5A). To interpret the biological significance of this HRV16-induced effect on splicing, we performed a pathway analysis using WebGestalt on the 1228 differentially spliced genes. While response to interferon was identified as a major pathway altered by HRV16 infection through alternative splicing, the functional enrichment analysis indicated that most differentially spliced genes during HRV16 infection were involved in the microtubule cytoskeleton required for cilia structure, ciliogenesis and cilia function (Fig. 5B).

**Figure 5.**
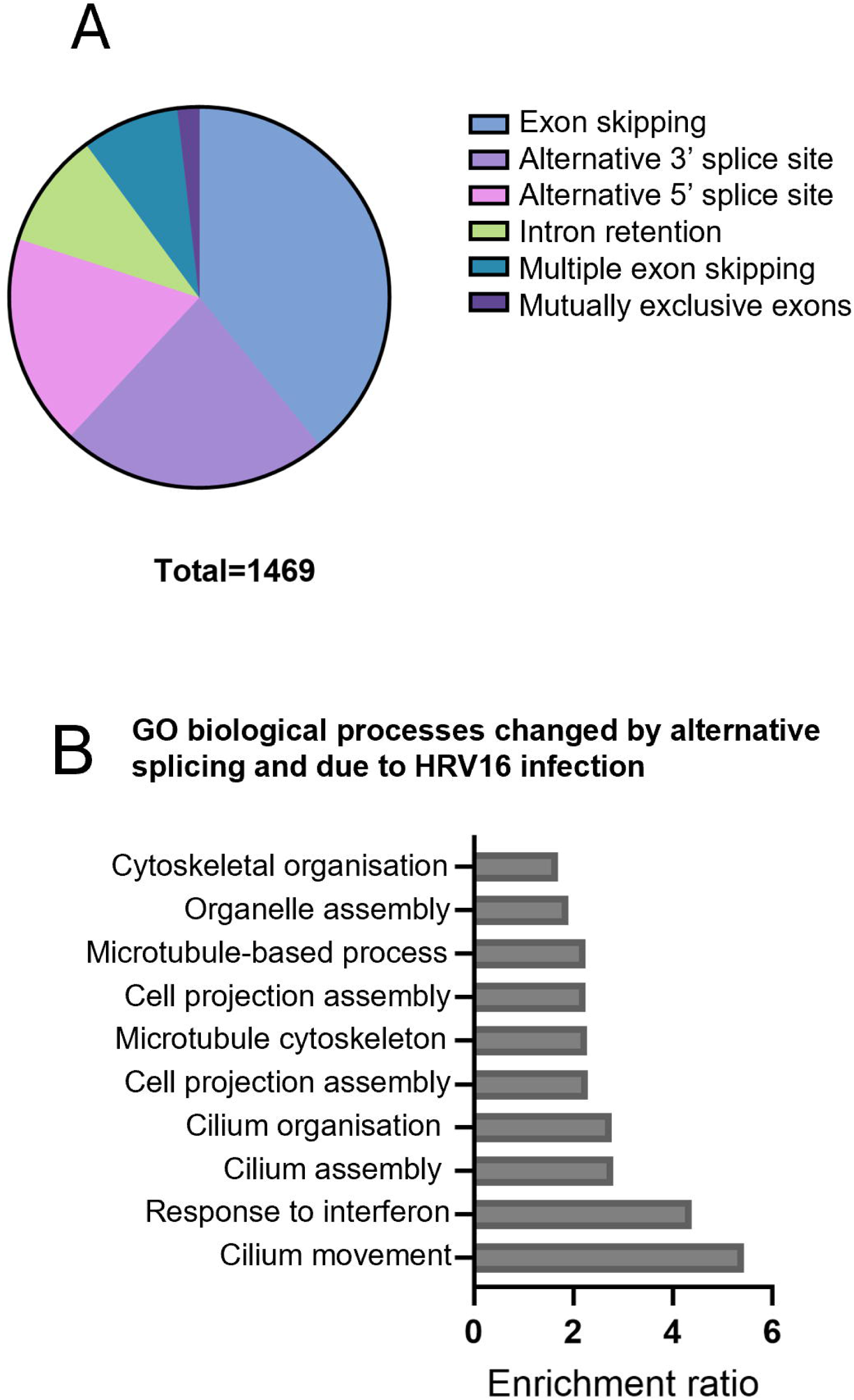
HRV16 infection in 3D cultured primary bronchial epithelial tissues leads to changes in the splicing of cilia-related genes. A. Pie chart of the relative proportions of the various splicing events identified in 1469 single alternative splicing events identified using SplAdder. B. Pathway analysis of the significantly differentially spliced genes during HRV16 infection in primary bronchial epithelial 3D cultures (p<0.05, student’s t-test, 2-tailed).

To verify specific alternative splicing changes due to HRV16 infection seen in the functional enrichment analysis we analysed three cilia-related genes whose alternative splicing is known to be related to ciliopathies [59]. These were radial spoke head protein 1 (RSPH1), which is located in the spokes of cilia and controls cilia motility [60]; intraflagellar transport protein 74 (IFT74), which is involved in protein transport within cilia and is required for ciliogenesis [61]; and transmembrane protein 67 (TMEM67), a ciliary transition zone protein required for cilia structure [62]. Figure 6A indicates the positions of these proteins in cilia. Sashimi plots showing the exon coverage from the RNA-Seq data were generated for the three selected genes using MISO (Fig. 6B-D). There was a small decrease in gene coverage for RSPH1 and TMEM67 indicating reduced transcription of the genes upon HRV16 infection. More importantly, clear differences in exon inclusion can be seen for each gene (see arrowheads indicated on the maps below the gene coverage profiles) comparing mock-infected to HRV16-infected cells suggesting that HRV16 infection alters mRNA splicing of these RNAs leading to mutations in the encoded proteins.

**Figure 6.**
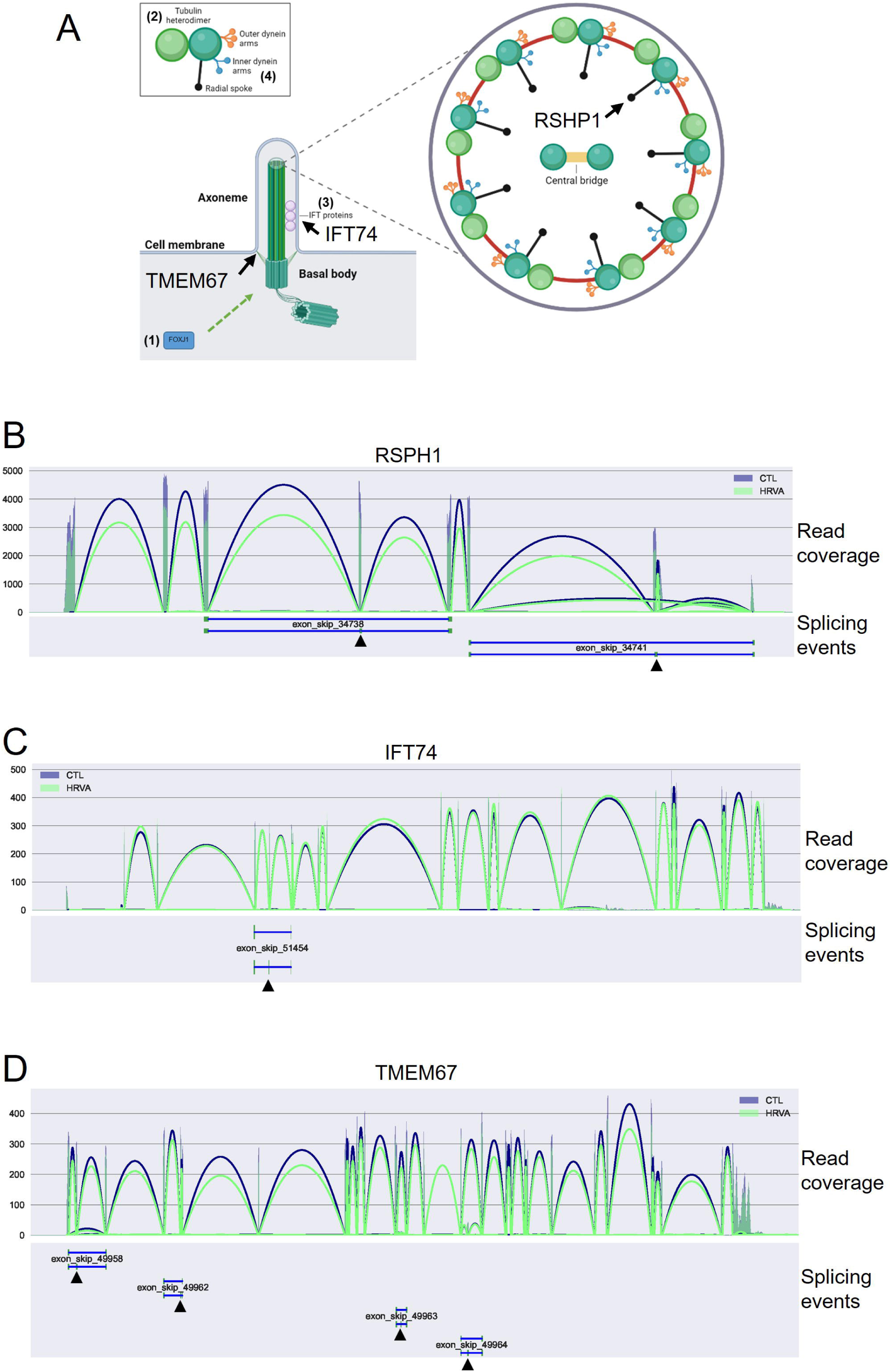
Representative changes in cilia-related RNA splicing using Sashimi plots. A. Diagram of the structure of a cilium showing the positions of the proteins encoded by the RNAs shown in B-D (generated using biorender: https://biorender.com). B-D. Sashimi plots of read coverage across genes encoding B. RSPH1, C. IFT74, and D. TMEM67. Dark blue lines, read coverage from mock-infected air-liquid interface cultures. Green lines, read coverage from HRV16-infected air-liquid interface cultures. Blue lines and black arrows beneath the Sashimi plots indicate skipped exons.

### HRV16 infection leads to impaired cilia in 3D cultured primary bronchial epithelial tissues

Based on the finding that HRV16 infection induced changes in the alternative splicing of genes involved in cilia development and function, we wanted to assess whether infection affected the numbers and morphology of cilia in our primary bronchial epithelial 3D model (Fig. 7). Figure 7A shows a representative image of an H&E stained, mock-infected 3D culture of HBECs. Figure 7B shows a representative image of an H&E stained HBEC tissue infected with HRV16 (3×10^6^ pfu) for up to 48 hours. HRV16-infected tissues showed reduced cilia density, and the cilia appeared shorter than those on mock-infected tissues (enlarged images in A and B). Cilia numbers were manually counted and, compared to mock infected tissues treated in exactly the same manner, infected tissues showing significantly decreased cilia numbers at all infection timepoints investigated (Fig. 7C). The largest decrease was recorded at 48 hours post infection, where infected samples had 4.5-fold less cilia compared to mock-infected samples. Additionally, the average cilium length in these respective samples was quantified (Fig. 7D). This analysis indicated that the average cilium length was significantly decreased in infected compared to mock-infected samples at 24 and 48 hours post infection. As with cilia numbers, the average cilium length showed the largest difference at 48 hpi, where the average cilium length in infected samples was decreased by 1.5-fold compared to mock-infected samples (Fig. 7D).

**Figure 7.**
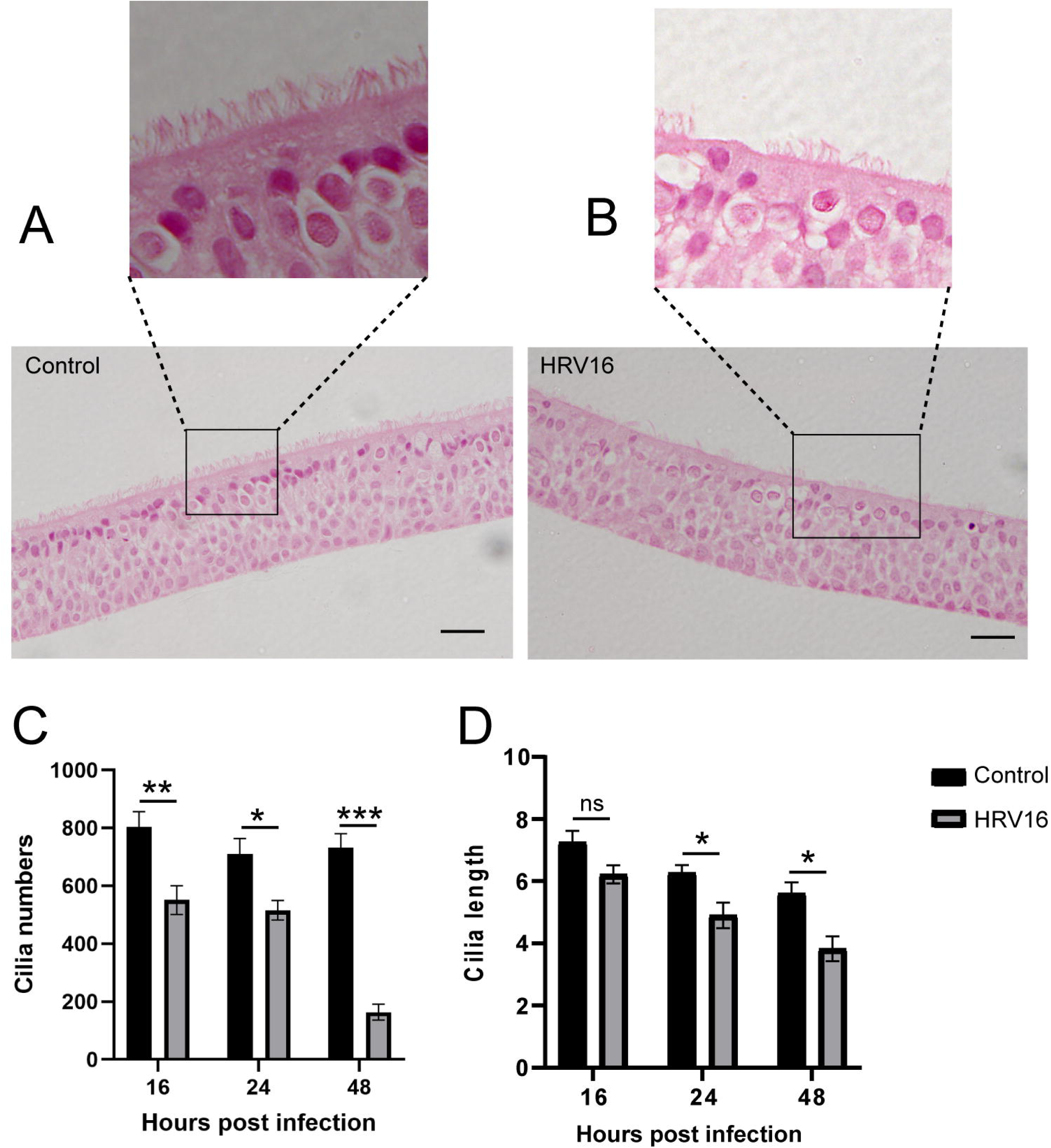
Cilia are reduced in number and in length due to HRV16 infection in 3D cultured primary bronchial epithelial tissues. A. H&E-stained 3D air-liquid interface culture grown for 4 weeks of mock-infected HBECs. A magnification of the upper surface of the tissue is shown above the main image. B. H&E-stained 3D air-liquid interface culture of HBECs grown for 4 weeks then infected with 3×10^6^ pfu HRV16 for 48 hours. Scale bars=50µm. A magnification of the upper surface of the tissue is shown above the main image. C. Graph of quantification of cilia numbers in mock-infected and HRV16-infected 3D cultures. D. Average cilium length on mock-infected and HRV16-infected 3D cutlures. Counts were taken from five sections from each of three replicate tissues at 16, 24 and 48 hours post-infection. The data show the mean and standard error of the mean. Significant statistical difference, *p=<0.05, **p,<0.005, ***p<0.0005 (students t-test, 2-tailed).

To confirm these data, we carried out immunofluorescence staining of sections of 3D cultures of HBECs either mock-infected or HRV16-infected. We chose to use antibodies against β-tubulin, a core cilium structural protein and TMEM67, a ciliary transition zone protein located at the junctions of cilia to the plasma membrane. β-tubulin was observed along the lengths of the cilia in mock-infected cells (Fig. 8 A, B) but levels of the protein were reduced in HRV16-infected cells and cilia were not clearly seen (Fig. 8 C, D). Quantification of β-tubulin staining showed a significant decrease at all time points, and this was especially significant at later times of infection (Fig. 8 I). TMEM67 staining for mock-infected tissues was found in cells in the mid layers and on the outer surface of the tissue, in discrete sub-cilia regions (Fig. 8 E, F). The location of staining was similar for HRV16-infected tissues, but the staining was more diffuse (Fig. 8 G, H). Quantification of staining showed a statistically significant decrease in TMEM67 levels in infected tissues at all time points (Fig. 8 J). Taken together these data suggest that HRV16 infection causes a reduction in cilia density and changes in cilia structure in the respiratory epithelium.

**Figure 8.**
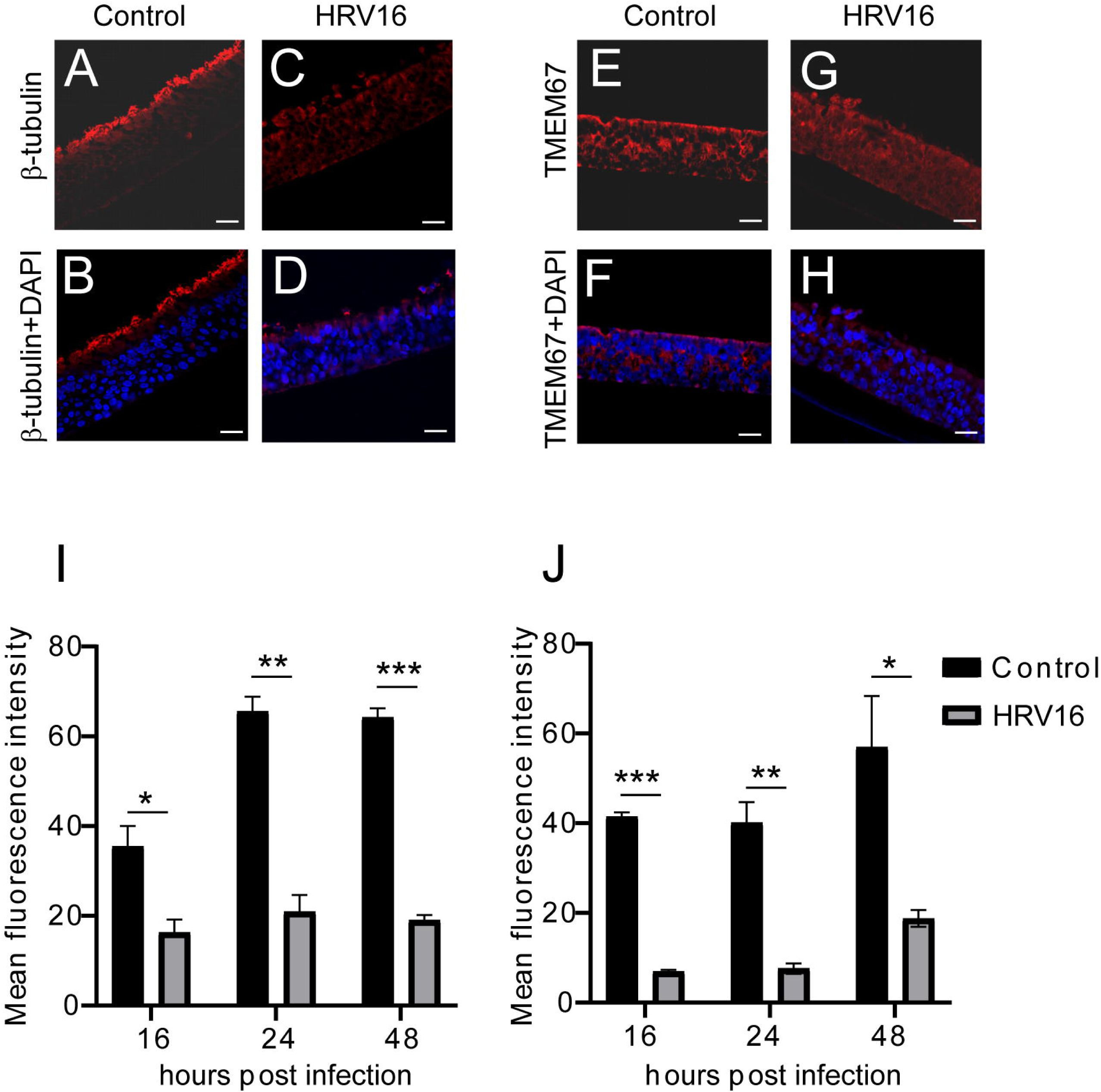
HRV16 infection of 3D cultured primary bronchial epithelial tissues leads to decreased expression of the cilia proteins β-tubulin and TMEM67. Fluorescence confocal microscopy image of HBEC 3D air-liquid interface cultures grown for 4 weeks using an antibody to detect β-tubulin in A. mock-infected (Control) and C. HRV16-infected (HRV16: 3×10^6^ pfu) tissues at 48 hours post-infection. B & D. The same images but with DAPI staining added to show the cell nuclei in the tissues. Fluorescence confocal microscopy image of an HBEC 3D air-liquid interface culture grown for 4 weeks using an antibody to detect TMEM67 in E. mock-infected (Control) and G. HRV16-infected (HRV16: 3×10^6^ pfu) tissues at 48 hours post-infection. F & H. The same images but with DAPI staining added to show the cell nuclei in the tissues. Scale bars=20µm. I. Graph of the quantification of staining intensity of the β-tubulin antibody over time in mock-infected (Control) and HRV16-infected (HRV16) 3D HBEC cultures. J. Graph of the quantification of staining intensity of the TMEM67 antibody over time in mock-infected (Control) and HRV16-infected (HRV16) 3D HBEC cultures. The data show the mean and standard error of the mean from 5 sections from three separate air-liquid interface cultures. *p<0.01, **p<0.001, ***p<0.0001 (student’s t-test, 2-tailed).

## Discussion

Human rhinovirus (HRV) is the most common respiratory virus infecting humans. Repeat infections in childhood can lead to wheezing and asthma, and these symptoms are exacerbated by subsequent HRV infections. In addition, in individuals with chronic obstructive pulmonary disease (COPD), HRV infection can lead to severe disease requiring hospitalisation. Discovering cellular factors that lead to exacerbations of HRV infection is key to designing novel therapies against these respiratory infections and conditions. This study has revealed two novel aspects of the cellular response to HRV16 infection. First, we have shown that the cellular splicing kinase SRPK1 is a restriction factor for HRV16 (a variant of HRV-A) infection in primary bronchial epithelial cells. Second, changes in alternative splicing cause by an HRV16 infection-induced down regulation of SRPK1 impact cilia structure and function in 3D cultured primary bronchial epithelial tissues.

Transcriptomics studies in nasal scrapings [7, 39], bronchial scrapings [63] and nasal and lung epithelial air-liquid interface 3D cultures [41, 43, 57] have shown that HRV16 infection alters expression of immunity-related genes. Our transcriptomic analysis of HBEC air-liquid interface cultures agrees with the conclusions of these studies. There was significant up-regulation of genes encoding proteins involved in anti-viral defence (e.g. OASL, MX2), and the interferon signalling pathway (e.g. IFNL1, INFL2) and several chemokines (e.g. CXCL10, CXCL11) known to be induced by HRV infection. Although SRPK1 is well known as a kinase that controls splicing via phosphorylation of SR proteins [29], it also positively regulates innate immunity to viral infections via NF-ĸB and interferon gamma [28]. Therefore, SRPK1 down-regulation during HRV16 infection could be beneficial to the infectious process, as we have demonstrated.

In an in vivo study of a mixed nasal cell population from children with wheeze infected with HRV, overall SRPK1 mRNA expression was upregulated compared to HRV-negative controls [7]. SRPK1 mRNA was also upregulated 1.3-fold after a 48 hour HRV16 infection of 3D HBEC tissues in this study. However, increased SRPK1 expression at the RNA level did not lead to increased levels of SRPK1 protein. In fact, in 3D HBEC tissues infected with HRV16 there was a significant decrease in SRPK1 levels and in phosphorylation of selected SR proteins compared to mock infected tissues. This result agrees with the decreased phosphorylation of SR proteins observed in a study where rhabdomyosarcoma cells were infected with another enterovirus, enterovirus A71 [64]. Our data suggest that HRV16 represses the activity of SRPK1 during infection. This could be either through decreased protein levels, as we have observed, or due to changes in cell signalling caused by HRV16 infection impacting SRPK1 kinase activity, which is itself controlled by Ck2/Akt-mediated phosphorylation [27].

However, inhibition of SRPK1 kinase activity has been shown to inhibit replication of viruses such as HIV, hepatitis C and Sindbis virus [65–68] and it has recently been reported that SRPK1 inhibition can reduced IRE-dependent translation and replication of enterovirus A71 [64]. These findings clearly suggest that SRPK1 is not a restriction factor of these viruses as it is required for their life cycles. In contrast, both inhibition and overexpression of SRPK1 has been shown to reduced Ebola viral replication [69], while SRPK1 is a known restriction factor for hepatitis B virus infection [70]. These diverse findings indicate that viruses may utilise SRPK1 in different ways for replication. Further, balancing activity of SRPK1 throughout virus infection may be key. For example, repression of SRPK1 stimulation of innate immunity early in infection could facilitate viral infection while upregulation later in infection could be required for viruses such as HIV and that require either cellular splicing [22] or enteroviruses that require SRPK1 activity for stimulation of viral RNA translation [64] to complete their life cycle.

Changes in expression of genes involved in cilia formation and function were found previously in a study comparing the transcriptome of air-liquid interface cultures established from the lungs of non-asthmatic versus asthmatic individuals [41] and changes were greater following HRV infection. Gene ontology analysis of RNA sequencing data showed that cilia function was found to be altered in human nasal epithelial cells infected with HRV16 [43]. Stimulation of the antiviral immune response by addition of poly(I:C) in human nasal epithelial stem/progenitor cells also revealed changes to genes involved in ciliogenesis and function [71]. In agreement with these studies, inspection of our DEG data from RNA sequencing showed reduced expression of genes encoding dynein axonemal assembly proteins, Bardet-Bield syndrome (BBS) genes important for cilia development and function, intraflagellar transport proteins, and tubulins, structural components of cilia.

The hypophosphorylation of SR proteins due to HRV16 infection in 3D HBEC tissues that we have observed would be expected to cause changes to cellular splicing [30, 72]. We observed a major impact on alternative splicing in HBEC tissues. Remarkably, when we analysed changes to gene expression due to altered splicing events, we discovered that most of the changes affected splicing of mRNAs encoding proteins involved in cilia structure and function. Interestingly, tissue-specific alternative splicing of cilia-related RNAs underlies a wide range of ciliopathies [59]. This suggests that correct alternative splicing is key to production of normal cilia proteins. Indeed, when we examined the morphology of the air-liquid interface cultures following HRV16 infection we found a blunting and loss of cilia compared to mock infected tissues. We propose that a combination of transcriptional down-regulation of cilia-related genes together with specific changes in splicing of cilia-related mRNAs could lead to malformed or damaged cilia. These morphological changes could lead to increased susceptibility to recurrent HRV infections and/or the development of chronic inflammatory disease.

There are several limitations to this study. Most importantly, the analyses were carried out using HBECs from a single donor. We cannot rule out the possibility that data from this donor may not reflect the general population. We studied virus infection of HBEC tissues at 48 hours post infection, and transcriptomic changes could be significantly different at other time points. As we only analysed transcriptomic changes in bronchial epithelial cells, a comparison of these changes between bronchial and nasal epithelial cells, grown as fully differentiated 3D tissues would have been interesting to perform. It is known that HRV infection causes redistribution to the cytoplasm of hnRNP proteins, known antagonists of SR proteins in alternative splicing [73], and ectopic expression of the HRV16 3C protease has also been shown to mislocate SRSF2, a splicing factor which we have not studied, in the nucleus of transfected cells. This means that changes in SR protein phosphorylation may only contribute to, rather than directly cause, the changes in splicing induced by HRV16 infection. Finally, examination of changes in expression of SRPK1, and its effects on its substrates, and any impact on ciliated epithelial cell function in HRV-C infected cells would be important to consider in the future since HRV-C infection can have more serious consequences, clinically.

## Conclusions

This study has revealed the cellular kinase SRPK1 as a new restriction factor for HRV16 infection in epithelial cells. The mechanism of restriction is not known. However, since SRPK1 can activate production of interferon, interferon response factors and certain chemokines [28], induction of innate immunity to viral infection seems likely. Indeed, transcriptomic analysis showed up-regulation by HRV16 of many innate immune factors. However, a key finding of this study is that HRV16 infection alters expression of splicing factors and their phosphorylation. Importantly, we found major changes in alternative splicing of RNAs encoding structural components of cilia. This suggests a molecular mechanism by which HRV infection might result in damage to the mucociliary compartment which would result in inefficient clearance of subsequent viral infection enabling infection to move to the lower airways, which would lead to exacerbations.

## Acknowledgements

This work was funded by a BBSRC Industrial Case CSV Training Award no. BB/R505341/1. Quan Gu is supported by an MRC award #MC_UU_00034/5. We are grateful to Chris McRae at Astra Zeneca for facilitating agreements between Astra Zeneca and the University of Glasgow and for arranging the industrial placement for student Chris Rozario (C.R.) in Astra Zeneca. We thank especially Jenny Horndahl, at AstraZeneca, Sweden for hosting C.R. during his industrial placement. We would like to thank the staff at the University of Glasgow’s Veterinary Diagnostics service for carrying out paraffin embedding, sectioning and staining of tissues. Prof Carl Goodyear, Director, the GLAZgo Discovery Centre helped arrange the studentship and provided critical feedback throughout the study.

## Declaration of Interests

Prof Maciewicz is retired from Astra Zeneca but at the start of the project she owned shared in Astra Zeneca. Astra Zeneca had no input into the design of the study or interpretation of the data.

## Author contributions

C.R. Investigation, formal analysis. writing, original draft

Q.G. Investigation, formal analysis, data curation, visualisation.

A.S. Data curation, resources.

R.M. Conceptualisation, supervision, funding acquisition, writing review and editing.

S.V.G. Conceptualisation, supervision, funding acquisition, visualisation, writing review and editing.

## Data availability statement

The underlying data for the graphs shown in this manuscript can be accessed at https://doi.org/10.5525/gla.researchdata.1946. The transcriptomic data can be found at RBI/ENA with the accession ID PRJEB88791.

## Notes

https://doi.org/10.5525/gla.researchdata.1946

