## Supplementary figures and images for "Human Rhinovirus 16 impacts cilia structure in 3D cultured primary bronchial epithelial tissue through alternative splicing of host cilia RNAs"

### Supplementary Figure1

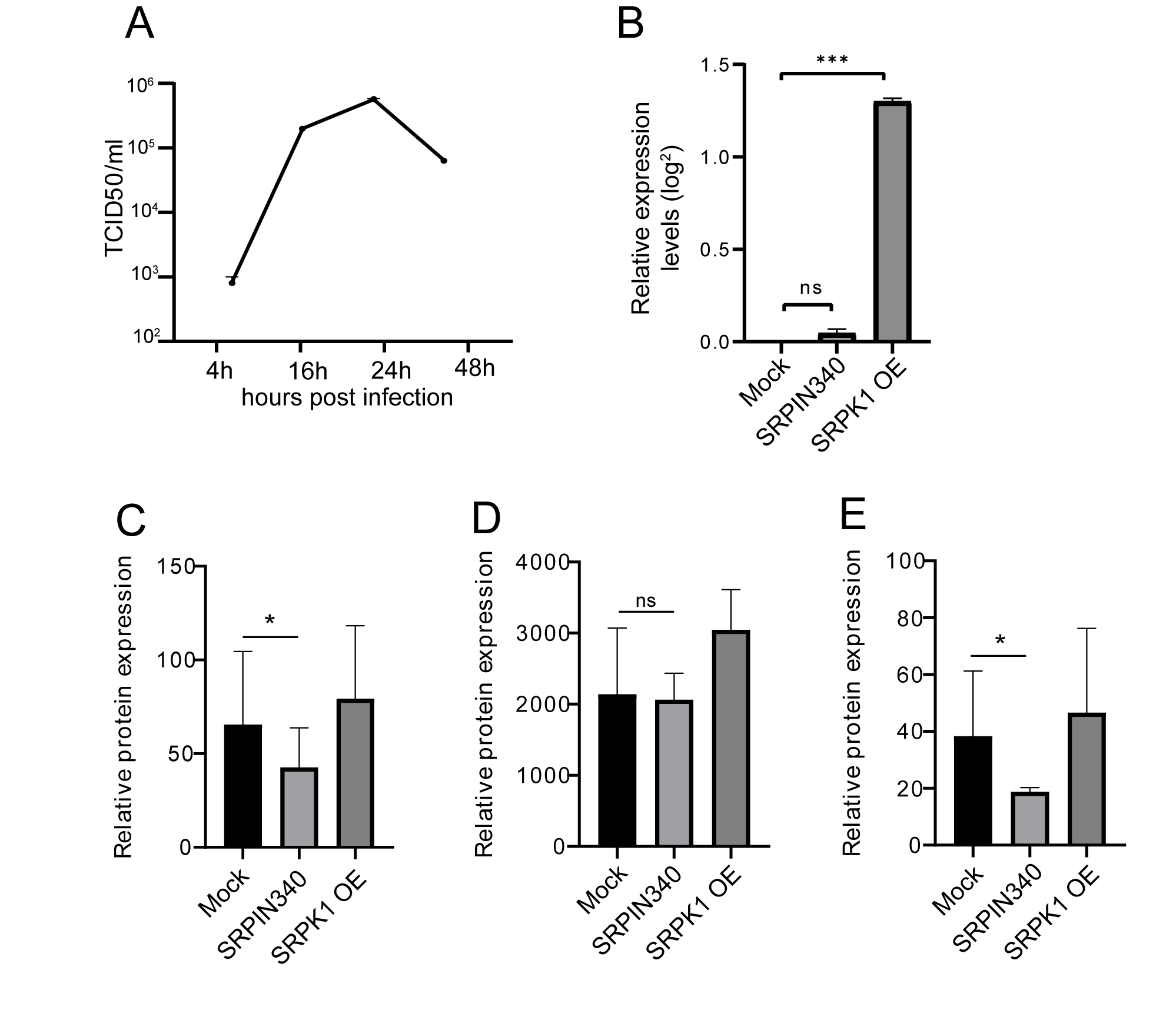

### Supplementary Figure2

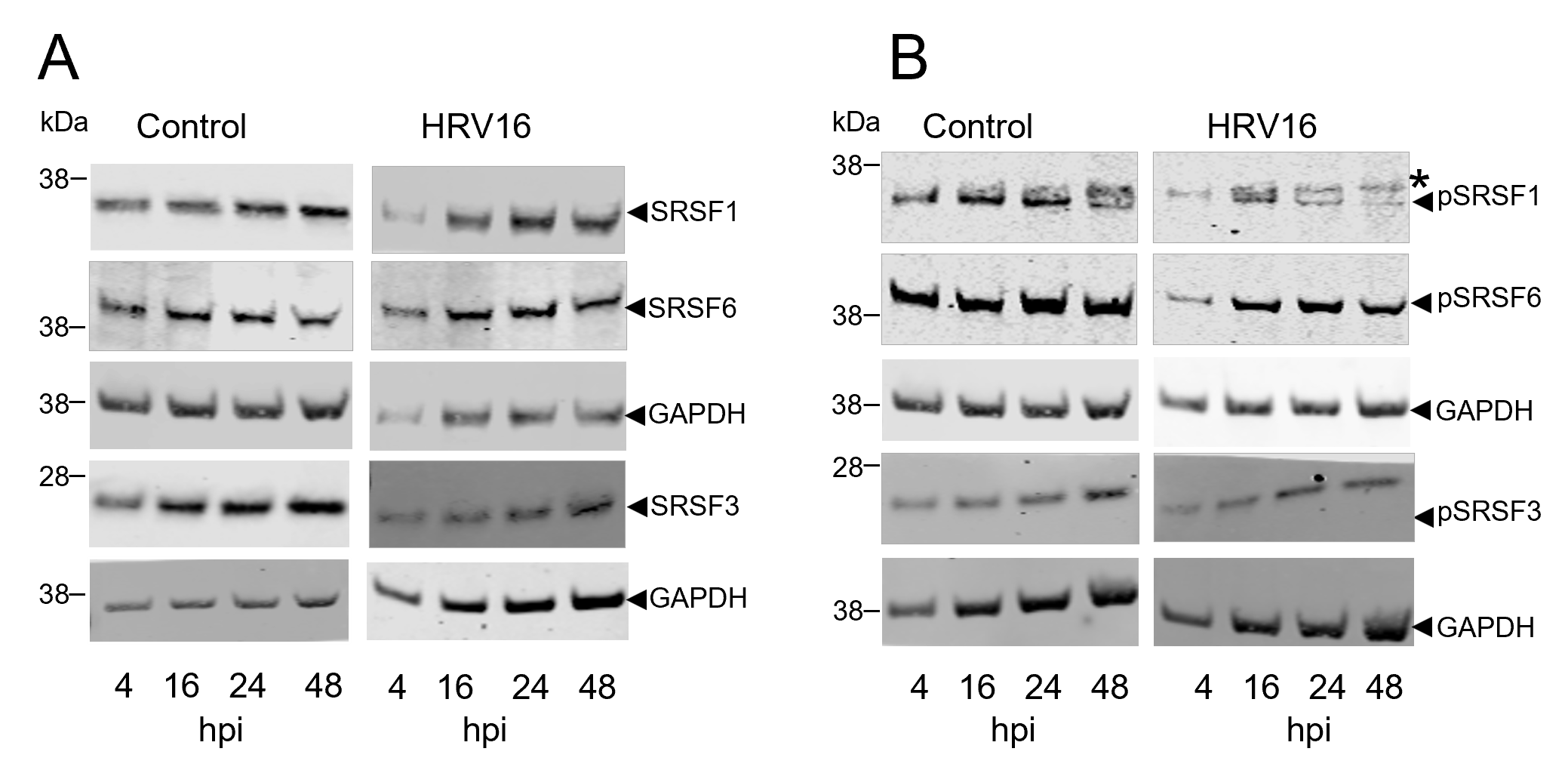

### Supplementary Figure 3

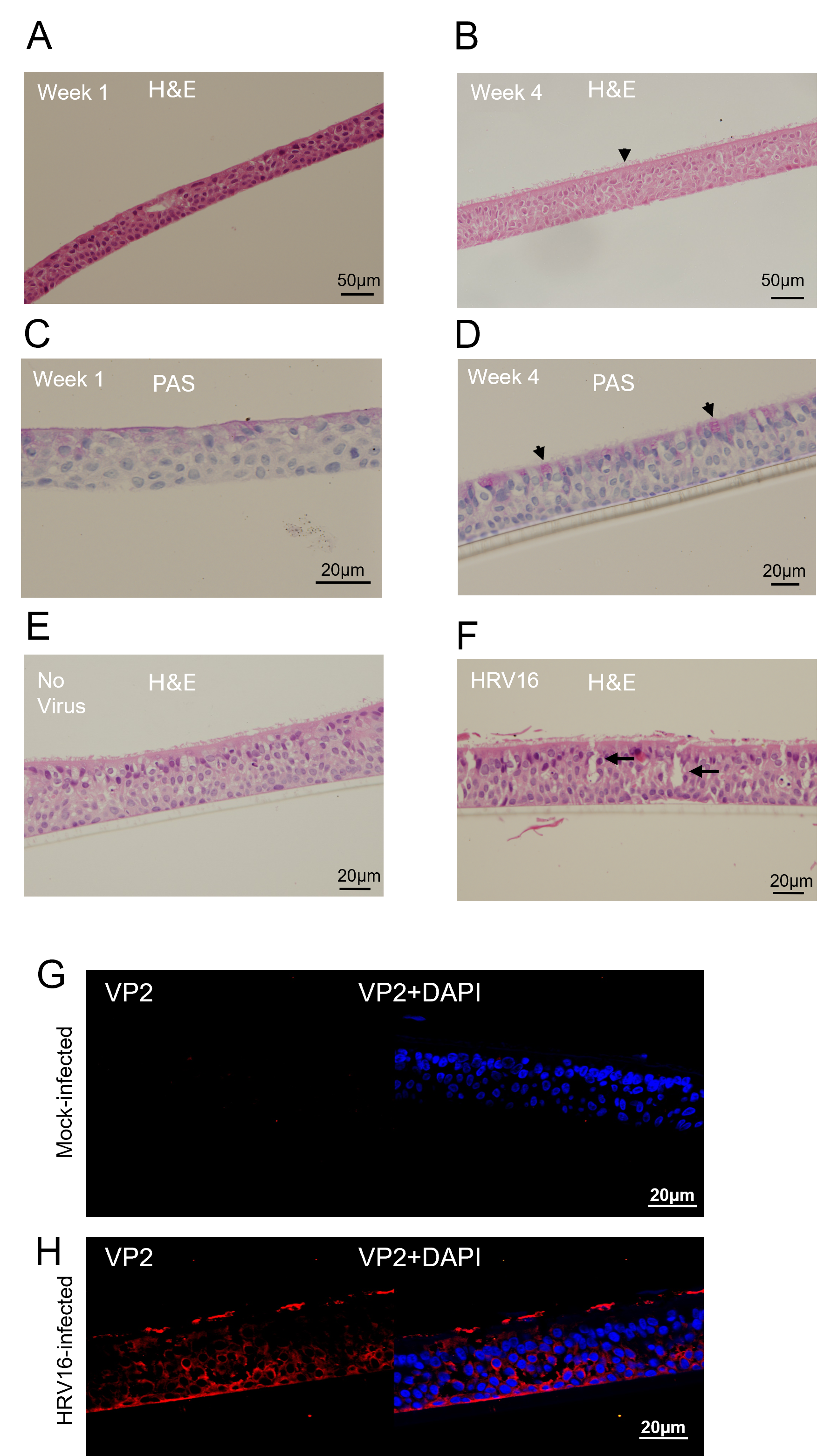

### Supplementary Figure 4

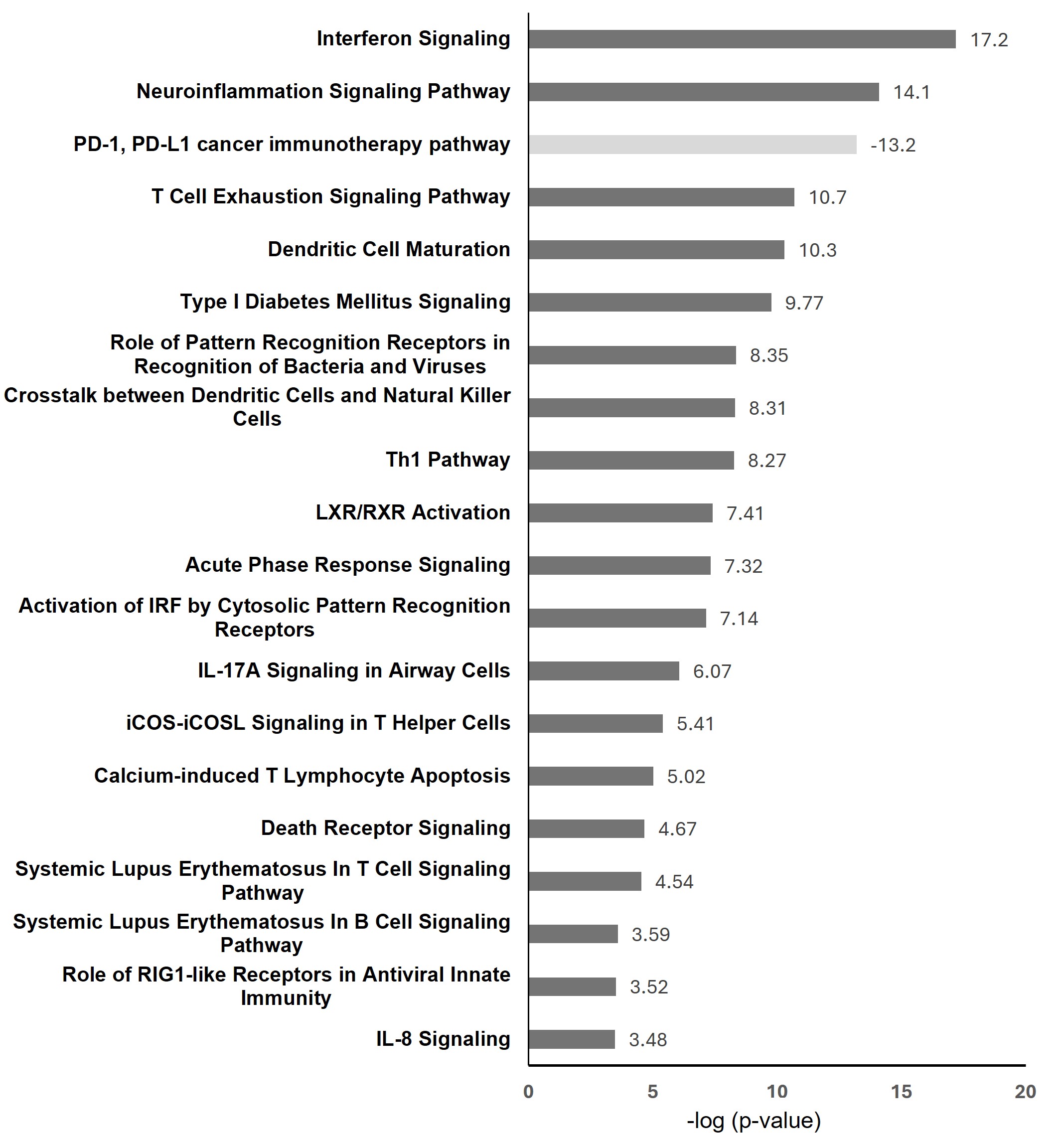
