## Supplementary data captions for "Human Rhinovirus 16 impacts cilia structure in 3D cultured primary bronchial epithelial tissue through alternative splicing of host cilia RNAs"

**Supporting information captions**

**Supplementary Figure 1. SRPIN340 treatment and SRPK1 overexpression alter phosphorylation of SR proteins in primary human bronchial epithelial cells (HBECs).** A. Graph of the time course of HRV16 infection (3x10^6^ pfu) in primary human bronchial epithelial cells. Error bars are not visible due to low level variation between replicates. B. Graph of relative mRNA expression levels of SRPK1 in HBECs DMSO-treated (Mock), SRPIN340-treated (SRPIN340) or transfected with an expression construct for SRPK1 (SRPK1 OE). Graph of relative levels of C. phosphorylated SRSF1, D. phosphorylated SRSF3, E. phosphorylated SRSF6 in HBECs DMSO-treated (Mock), SRPIN340-treated (SRPIN340) or transfected with an expression construct for SRPK1 (SRPK1 OE). The graphs show the mean and standard error of the mean from three separate experiments. ns, no significant difference, * p>0.05, *** p>0.0005 (student’s t-test).

**Supplementary Figure 2. Western blots showing representative primary data used for the values shown in the graphs in Figure 2.** Control, mock infected HBECs. HRV16, HBECs infected with HRV16 at an MOI=3 (hpi, hours post infection). The asterisk indicates another SR protein, SRSF2, which is a poor substrate of SRPK1, which has a larger mass than SRSF1.

**Supplementary Figure 3. HRV16 infection changes the morphology of human bronchial epithelial cells (HBECs) in 3D cultures.** A. A representative HBEC tissue grown in air-liquid interface culture for one week and H&E stained. B. A representative HBEC tissue grown in air-liquid interface culture for four weeks and H&E stained. The arrowhead indicates cilia on the upper surface of the tissue. Scale bars = 50µm. C. PAS staining (pink) of an HBEC tissue grown in air-liquid interface culture for one week. D. PAS staining (pink) of an HBEC tissue grown in air-liquid interface culture for four weeks. Arrowheads indicate goblet cells. Tissues are counterstained with haematoxylin (blue) to show the nuclei. Scale bars = 20µm. E. H&E stained representative HBEC tissue grown in air-liquid interface culture for four weeks and mock infected. F. H&E stained representative HBEC tissue grown in air-liquid interface culture for four weeks and infected with 3x10^6^ pfu HRV16 for 48 hours. Arrows indicate mechanical damage. Scale bars = 20µm. G. Immunofluoresence microscopy of a section of HBEC tissue grown in air-liquid interface culture for four weeks, mock infected and stained with an antibody against VP2 (red). H. Immunofluoresence microscopy of a section of HBEC tissue grown in air-liquid interface culture for four weeks, infected with 3x10^6^ pfu HRV16 for 48 hours and stained with an antibody against VP2 (red). The images on the right hand side of the panels in G and H show staining with DAPI (blue). Scale bars = 20µm.

**Supplementary Figure 4. Pathway analysis of differentially expressed genes comparing mock-infected to HRV16-infected bronchial epithelial cells 3D cultures.** The top twenty pathways enriched following IPA analysis of differentially expressed genes with a log2-fold change of >1 and <-1 are shown. Black bars indicate upregualted pathways. Grey bar represents a downregulated pathway. Numbers after the bars indicate levels of up-regulation (or down-regulation: -ve value, grey bar) of each factor.

**Supplementary Table 1. List of all 4034 DEGs due to HRV16 infection** (false discovery rate (FDR)<0.05, Benjamini-Hochberg procedure).

**Supplementary Table 2. List of DEGs selected for over-representation analysis to identify cellular pathways altered by HRV16 infection.** The subset of 655 DEGs selected for pathway analysis, with log2-fold change values within the range of >1 to <-1
